# Pulmonary maternal immune activation does not extend through the placenta but leads to fetal metabolic adaptation

**DOI:** 10.1101/2023.03.17.532052

**Authors:** Signe Schmidt Kjølner Hansen, Robert Krautz, Daria Rago, Jesper Havelund, Nils J. Færgeman, Audrey Prézelin, Julie Rivière, Anne Couturier-Tarrade, Vyacheslav Akimov, Blagoy Blagoev, Betina Elfving, Arnaud Stigliani, Ulla Birgitte Vogel, Konstantin Khodosevich, Karin Sørig Hougaard, Albin Sandelin

## Abstract

Maternal immune system activation (MIA) during pregnancy can disrupt the fetal environment, causing postnatal susceptibility to disorders. How the placenta and the fetus respond to acute MIA over time is unknown. Here, we characterized the response to acute maternal pulmonary inflammation across time in maternal and fetal organs using multi-omics. Unlike maternal organs which mounted strong innate immune responses, the placenta upregulated tissue-integrity genes, likely to prevent fetal exposure to infections, and downregulated growth-associated genes. Subsequently, the placenta upregulated biosynthesis and endoplasmic reticulum stress genes in order to return to homeostasis. These responses likely protected the fetus, since we observed no immune response in fetal liver. Instead, likely due to nutrient depletion, the fetal liver displayed metabolic adaptations, including increases in lipids containing docosahexaenoic acid, crucial for fetal brain development. Our study shows, for the first time, the integrated temporal response to pulmonary MIA across maternal and fetal organs.

## INTRODUCTION

Activation of the maternal immune system during pregnancy poses a risk for the developing fetus. Long-term implications of maternal immune activation (MIA) include, but are not limited to, changes in offspring metabolic and immune function and predisposition to neurodevelopmental disorders^1–7^. As the anatomical interface between mother and fetus, the placenta has a remarkable range of physiological functions that are pivotal for fetal development, e.g. transfer of nutrients, efflux of waste, immunological protection and production of hormones^8^. Nutrient and oxygen transfer is contingent on its rich vasculature and permeability, which makes it receptive to maternal blood-borne pathogens and inflammatory mediators.

Studies have assessed the acute manifestation of maternal intraperitoneal or intrauterine inflammation in maternal organs, the placenta or fetal organs following exposure to LPS, vira or poly i:C^9,10^. Placental responses include leukocyte infiltration^11^, trophoblastic hypertrophy, altered mitochondrial function and energy metabolism and vascular reactivity^12^. Placental cytokine signaling was found to be important for fetal neurodevelopmental phenotypes^13^.

Less is known about short-term placental and fetal manifestations of MIA when the immunogen is targeted to the lungs, as in respiratory infections and maternal airway inflammation^14–20^, which are common during pregnancy: 49.6% of control mothers in the National Birth Defects Prevention Study reported respiratory infections during pregnancy^21,22^.

A limitation of many MIA studies is that they typically only measure single responsible mediators and/or capture the response in a single organ at a single time point. We argue that to understand the consequences of MIA and the fetal impact, characterization of changes over time in both maternal and fetal organs, including the placenta, is necessary.

To this end, we here characterize the temporal response to acute maternal lung inflammation across maternal and fetal organs, using multi-omics, imaging and integrative analyses. We show that maternal organs responded strongly, whereas the placenta orchestrated a specific adaptive response, comprising decreased cell growth and increased tissue integrity. Although this protected the fetal liver from an inflammatory response, it reacted to altered nutrient levels by metabolic adaptations, including production of lipids containing docosahexaenoic acid, crucial for fetal brain development.

## RESULTS

### Strong acute-phase pulmonary response to LPS does not extend to the placenta

To investigate gene expression response following MIA, we exposed pregnant C57BL/6 mice (gestation day 17) via the airways to 1 µg of lipopolysaccharide (LPS) or H_2_O vehicle (Ctrl) by intratracheal instillation: see Note S1 and Fig.S1A-B for dose choice considerations.

Mice were sacrificed 2, 5, 12 or 24h after instillation. At each time point, maternal lung and liver, placenta, and fetal liver were excised (Fig.1A). Fetal tail DNA was genotyped for sex and only one female pup per dam was used for each assay. The placenta (chorionic plate, labyrinth and junctional zones) and decidua were separated manually (Fig.1A). RNA was extracted from 7-9 biological replicates for each combination of treatment, time point and tissue (total 370 samples, from 74 dam-fetus pairs). Lung mRNA levels of the acute phase Serum amyloid A gene (*Saa3*)^23^, measured by qPCR, were highly increased throughout the 24h in LPS compared to Ctrl lungs (Fig.S1C). Pulmonary *Saa3* mRNA expression correlates closely with neutrophil influx after LPS exposure^24^. Hence, LPS caused strong pulmonary inflammation, consistent with^25,26^. LPS exposure induced weight loss, plateauing at 12 hours at 6-8% decrease vs. time-matched Ctrl (Fig.S1D)

**Figure 1:**
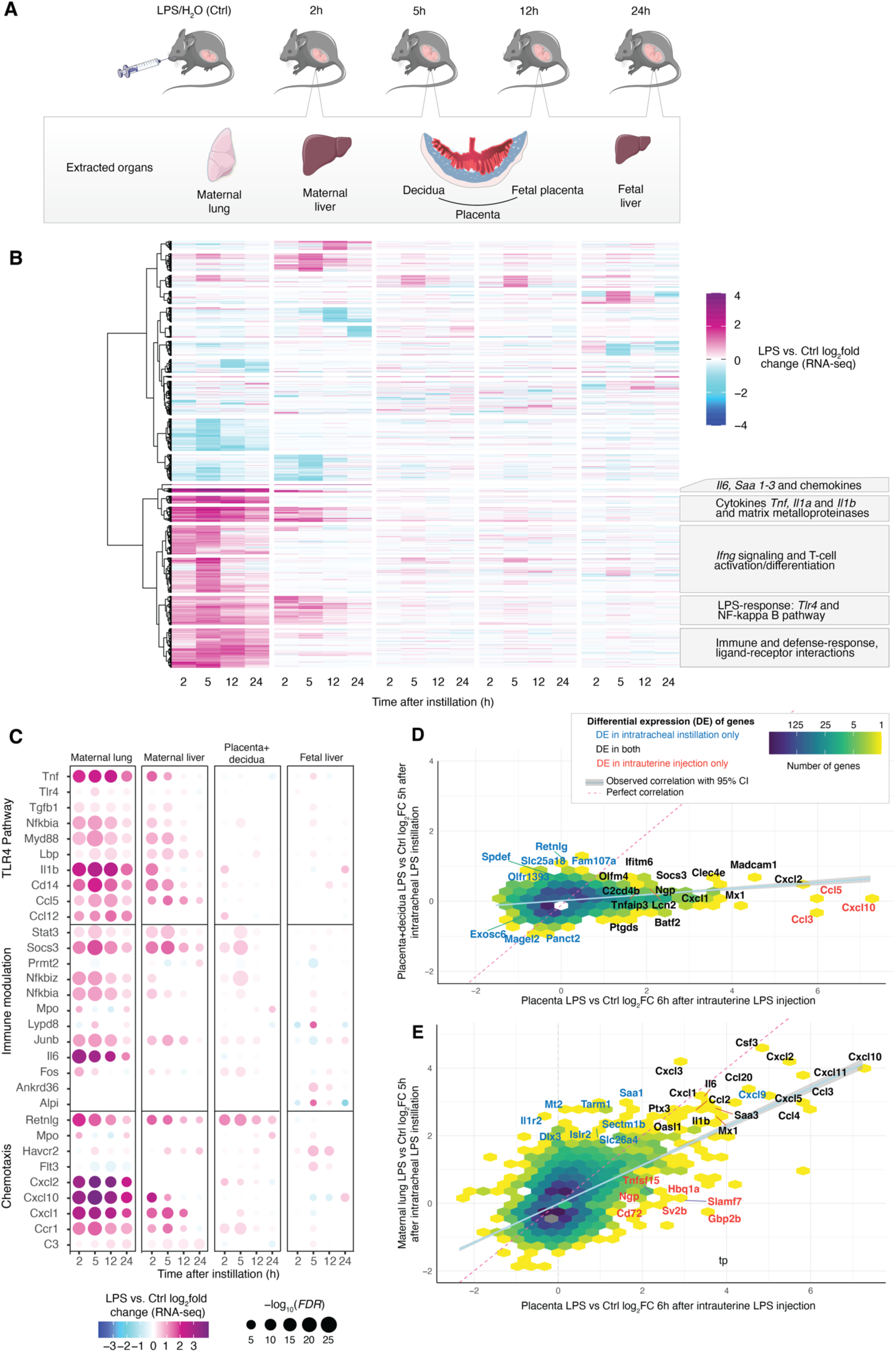
Experimental design and overview of RNA-seq data. **A: Experimental design.** At gestation day 17, C57BL/6 mice were instilled with either LPS or vehicle (H_2_0, denoted Ctrl). At set time points after instillation, mice were sacrificed and organs harvested (gray box) from mother and female fetuses. For each combination of treatment and time point, 7-9 mice were sacrificed. These organs were used for subsequent analyses, including RNA-seq (panel B). Blood plasma samples were also collected from mothers at all time points (not shown). **B: Overview of RNA-seq results.** Heat map rows show genes that changed expression ≥2-fold in at least one time point and tissue. Columns show time points after LPS or Ctrl instillation, sorted first by tissue (as indicated by gray box from panel A) and then by time (X axis). Colors indicate LPS vs Ctrl gene expression log_2_ fold change (log_2_FC), for respective time point and tissue. Callouts show major enrichments of GO terms for gene expression subclusters dominated by maternal lung and liver. **C: Expression change of selected genes and pathways.** The heat map is organized as panel B, but uses placenta+decidua samples for DE analysis and shows both expression change (placenta+decidua log_2_FC, indicated by color) and statistical significance (-log_10_*FDR*), indicated by dot size. Genes are grouped by pathway or functional role, as indicated on the right. **D: Comparison of placental gene expression change following intratracheal instillation vs. intrauterine LPS injection.** The X-axis shows placental gene expression log_2_FC of LPS vs Ctrl 6h after intrauterine LPS injection (data from^12^). The Y-axis shows placenta+decidua gene expression log_2_FC, LPS vs Ctrl, 5h after intratracheal LPS instillation (our data). Hexagon colors indicate the number of genes in the same area of the plot. Callouts show differentially expressed genes, colored by whether the gene was differentially expressed in one or both experiments. Dotted line indicates perfect correlation between X and Y: blue line indicates the observed correlation with 95% confidence interval in gray. **E: Comparison of maternal lung gene expression change following intratracheal LPS instillation vs. placental gene expression following intrauterine LPS injection.** The plot is organized as in panel D, but Y axis shows maternal lung gene expression log_2_FC of LPS vs Ctrl 5h after intratracheal LPS instillation (our data).

RNA samples were subjected to paired-end, polyA-selected RNA-seq were mapped to the mouse transcriptome (M23); all 370 libraries were retained after quality control. Principal component analysis showed that samples clustered by tissue and time point (Fig.S1E,F). We calculated the average LPS vs. Ctrl log_2_ expression fold change (log_2_FC) for each tissue and time point and visualized genes with an absolute log_2_FC>1 in at least one tissue and time point as a heatmap (Fig.1B). This enabled four important observations, also confirmed by differential expression (DE) analysis (Fig.S1G):

First, the maternal lung showed strong transcriptional responses at 2-5h, partially persisting at 12-24h, consistent with^25,26^. Gene ontology (GO) analysis showed that upregulated biological processes were, as expected^25–27^, dominated by pathways related to acute-phase signaling including response to lipopolysaccharide (e.g. TLR4 pathway), cytokine production, neutrophil migration and fever generation (Fig.1B, S1H).

Second, the maternal liver responded to LPS, albeit less strongly. The main response occurred at 2-5h. Many upregulated genes were shared with the lung and related to activation of innate immune response and inflammatory pathways, indicating a direct response to LPS^28^ (Fig.1B, S1I).

Third, the placenta and decidua also responded to LPS, but less strongly and qualitatively different from maternal lung and liver. Genes responded mainly at a single time point: the largest change occurred at 5h (Fig.1B). Notably, decidual and placental LPS vs Ctrl responses were highly similar; only a handful of genes showed a significant response in only one of the tissues (Fig.1B, S1J). This may reflect similar response patterns, or the difficulty in separating tissues of maternal and fetal origin^29^. Therefore, when analyzing expression response to LPS (e.g by LPS vs Ctrl log_2_FC), like in DE analysis, we combined placental and decidual samples, and denoted this ‘placenta+decidua’. For clarity, when analyzing gene expression levels in respective tissue rather than LPS response (e.g. transcripts per million (TPM)), we use ‘decidua’ and ‘placenta’ expression in the text.

Fourth, the fetal liver displayed a unique response to LPS compared to other tissues, but with a magnitude similar to that of the placenta in terms of number of DE genes (Fig.S1G) and range of log_2_FC values. As in placenta, the largest number of DE genes was observed at 5h.

Thus, there was a stark contrast between the strong and long-lasting LPS response in maternal lung and liver and the fainter and temporally more restricted response in placenta, decidua and fetal liver. As the placenta regulates maternal-fetal signaling^30^, its response is key to understanding placental and fetal strategies to ameliorate the consequences of maternal systemic inflammation. Thus, we profiled expression of key immune-related genes in placenta+decidua as a proxy for potential immune response and compared these across organs, starting with the canonical LPS-TLR4 pathway.

Although placental and decidual cells can induce the TLR4-pathway upon LPS stimulation^31–35^, we observed little or no LPS response in this pathway in placenta+decidua at any time point (Fig.1C, top). We speculated whether the placenta responded indirectly to LPS. Indeed, the expression of two groups of immune-related genes changed the most in placenta+decidua at 5h (Fig.1C, middle). Some were involved in signaling downstream of the TLR4-pathway, including *Nfkbiz*, an activator of Il6 but inhibitor of NFKB and Tnf signaling^36^. Also, immuno-modulatory genes of the IL-6/JAK/STAT pathway were upregulated, including *Stat3*, *Socs3, Junb* and *Fos*. Importantly, Il6 is one of the most highly induced cytokines by the NFKB pathway upon binding of LPS to TLR4^37^, and its inflammatory functions are inhibited by SOCS3^38,39^. Notably, *Il6* was highly induced in maternal lung only (Fig.1C, left panel). Only few chemoattractant genes were induced in placenta+decidua, including *Cxcl1*, *Cxcl2* and *Retnlg* (Fig.1C, bottom), suggesting negligible immune-cell infiltration.

To compare this presumably indirect response with direct placental LPS response, we compared our 5h placenta+decidua RNA-seq data to RNA-seq data following intrauterine LPS injection from^12^. The datasets were comparable in terms of day of gestation (GD17.5 and GD17) and time after exposure (5h and 6h), although the LPS dose (50 ug) aimed at inducing preterm birth (at approx. 12h after exposure) and is considerably higher than ours (1 ug). The placenta+decidua responses to intratracheal LPS-instillation (our data) and intrauterine LPS-injection^12^ were moderately correlated: a small set of genes were upregulated in both experiments, including antimicrobial and immune effectors, e.g. *Socs3, Clec4e, Batf2, Lcn2, Madcam1, Olfm4* and *Mx1* (Fig.1D). Intratracheal LPS-instillation, but not intrauterine LPS-injection, induced upregulation of a diverse set of genes including actin-dynamics modulator *Fam107a,* and immunomodulator *Retnlg*. Conversely, only intrauterine LPS-injection highly upregulated the chemokines *Ccl3, Ccl5 and Cxcl10* (Fig.1D). Strikingly, the placental expression changes following intrauterine LPS-injection at 6h were highly correlated to those of the LPS-instilled lungs at 5h in our experiment (Fig.1E), irrespective of the large difference in dose. We conclude that although the placenta has the capacity to mount a strong direct LPS response^12,34,40^, possibly contingent on direct, high LPS exposure, it reacts fundamentally differently to LPS instilled into maternal lungs, characterized by immune modulation rather than activation.

### Circulating Il6 is a candidate for inducing placental immune-modulation

We hypothesized that the placental gene expression response was contingent on one or more circulating cytokines/chemokines secreted by the maternal lungs in response to LPS^41^. Therefore, we measured concentrations of 11 cyto- and chemokines (Ccl2, 4, 7, 11, 17, 20, 24, Cxcl1, 13, 16 and Il6) in maternal plasma at all time points using the Luminex platform. We compared maternal plasma protein abundance changes (LPS vs. Ctrl log_2_FC by Luminex) with expression changes of the corresponding mRNA in maternal lung (LPS vs. Ctrl log_2_FC by RNA-seq; Fig 2A). Generally, cyto/chemokine concentrations in maternal blood were increased at 2h but decreased to Ctrl levels at 24h (Fig.2A). Corresponding lung mRNA levels were generally increased at 2h but decreased slower than protein levels over time. Notably, *Il6* and *Ccl20* mRNAs were highly upregulated in LPS-instilled lungs, especially at 2, 5 and 12h (log_2_FC=2.8-3.5). Their maternal blood protein levels were highly increased at 2h (log_2_FC>1.5) and then gradually decreased, reaching Ctrl-levels at 24h (Fig.5A, left). The levels of other measured chemokines fell into two categories: i) Ccl7, Ccl11 and Cxcl13 were 2-fold upregulated in blood at 2h (Fig.2A, middle) but generally not as highly induced at mRNA levels in lung, and ii) Cxcl1, Ccl2, Ccl4, Ccl17, Ccl16 and Ccl24, which were only modestly upregulated in blood (log_2_FC<1 at all time points), even though *Cxcl1, Ccl2 and Ccl4* mRNAs were highly upregulated in lung (Fig.2A, right).

**Figure 2:**
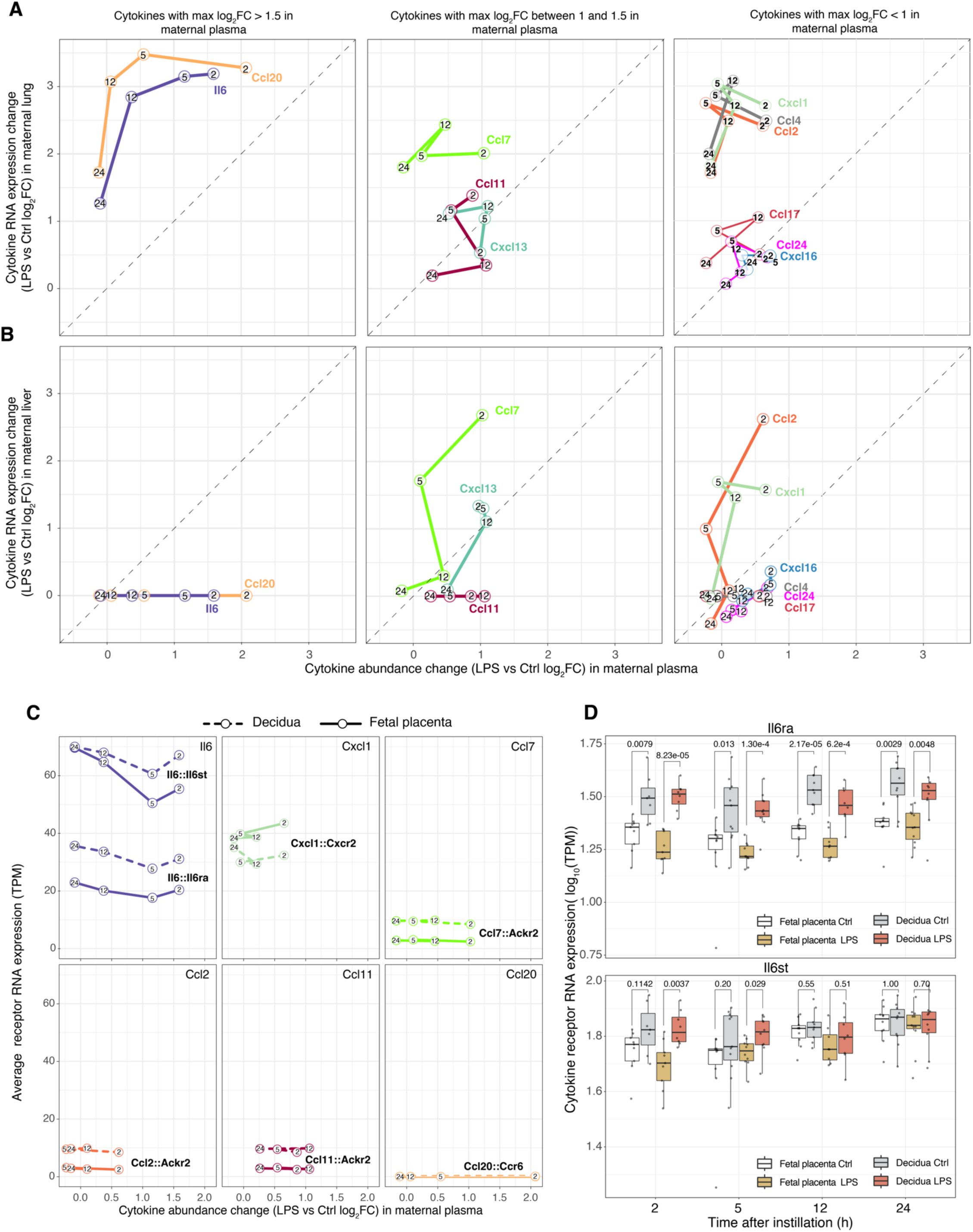
Exploration of signaling ligands and placenta cognate receptors. **A: Comparison of cytokine mRNA expression in maternal lung and protein levels in maternal plasma.** The Y axis shows mRNA expression change (LPS vs. Ctrl log_2_FC) of selected cytokines based on RNA-seq from maternal lung. The X axis shows abundance change (LPS vs Ctrl log_2_FC) of the corresponding cytokine proteins in maternal plasma, based on Luminex data. Each cytokine is indicated by color: numbers in connected dots indicate time after lung instillation. Each panel shows sets of cytokines in a specified log_2_FC range in maternal plasma as indicated on top. Dotted lines show perfect correlations (X=Y). **B: Comparison of cytokine mRNA expression in maternal liver and peptide levels in maternal plasma.** The plot is arranged as in A, but the Y axis shows mRNA expression change (LPS vs. Ctrl log_2_FC) of selected cytokines based on RNA-seq from maternal liver. **C: Comparison of cyto/chemokine abundance change in maternal plasma vs. cytokine receptor mRNA expression in decidua and fetal placenta.** Each subplot compares the abundance change (LPS vs Ctrl log_2_FC) of a selected cyto/chemokine (indicated on top) protein in maternal plasma (as in panel A) and the average mRNA expression level of annotated receptor(s) of this cyto/chemokine in Ctrl decidua (dotted lines) or placenta (solid lines), as RNA-seq TPM. Numbers in dots indicate time after lung instillation. Cytokine-receptor pairs are indicated as ‘cytokine::receptor’. Subplots are arranged by expression of their cognate receptor(s). Colors indicate cyto/chemokine, using the same color scheme as in panels A+B. **C: mRNA expression levels of Il6 receptors in decidua and fetal placenta** The Y axis shows mRNA expression distributions of Il6 receptors *Il6ra* and *Il6rst* as log_10_(TPM). Each dot corresponds to one sample. The X axis shows time after exposure. Colors indicate exposure and tissue type. *P* values of two-sided Mann-Whitney tests between relevant distributions are shown on top.

Therefore, we reasoned that Il6 and Ccl20 were most apparent candidates while Ccl7, Ccl11 and Cxcl13 might mediate local inflammatory responses. Because the liver contributes to combating systemic inflammation, it might constitute an additional source of cyto/chemokines^42^. Therefore, we also compared upregulation of cyto/chemokines in maternal blood with mRNA upregulation in maternal liver (Fig.2B). Strikingly, neither *Il6* nor *Ccl20* expression was upregulated in maternal liver (Fig.2B, left), although *Ccl7* and *Cxcl13* expression was moderately increased (Fig.2B, middle). Among the remaining measured cytokines which were not highly upregulated in blood, only *Ccl2* and *Cxcl1* mRNAs were highly upregulated in the liver (Fig.5B, right). Therefore, while maternal liver might contribute to the temporary increase in Ccl7 and Cxcl13 blood levels, the prominent increase in Il6 and Ccl20 in maternal blood is likely due to their high mRNA upregulation in the lungs.

We reasoned that the chemo/cytokines most likely inducing placental responses would have increased levels in maternal blood and their cognate receptor(s) would be highly expressed in the placenta. Hence, we correlated changes in cyto/chemokine blood concentrations (Luminex log_2_FC, LPS vs Ctrl) with average mRNA expression levels (RNA-seq TPM) of their annotated receptor(s)^43^ in placenta and decidua. Since receptors must be present prior to MIA to elicit effects, we used receptor expressions in Ctrl.

IL-6 was the most prominent candidate: its levels were highly increased in maternal blood, and its two receptor-subunits, *Il6ra* and *Il6st*, were highly and constitutively expressed in decidua and placenta (Fig.2C, top left), consistent with that IL-6 is a known mediator of MIA^2,13,44,45^. Interestingly, decidua had higher expression levels of both receptor-subunits than placenta regardless of treatment (significant at all time points for *Il6ra*, but only at some time points for *Il6st* (*P*<0.05, two-sided Whitney-Mann tests), suggesting that IL-6 receptor binding and downstream signaling may occur more in decidua than placenta (Fig.5C-D).

The Cxcl1 receptor *Cxcr2* was highly expressed in Ctrl decidua and placenta at all time points, while LPS increased Cxcl1 levels in maternal blood at 2h, suggesting a potential early interaction (Fig.2C, top middle). Notably, Cxcl1-Cxcr2 signaling in epithelial cells maintains immune-tolerance during pregnancy^46^.

The Atypical chemokine receptor 2 (*Ackr2*) was consistently expressed at low levels in decidua and placenta (Fig.2C,top tight and bottom left). It scavenges inflammatory CC chemokines, including Ccl2, 7 and 11, which are chemoattractants for immune cells^47,48^. Since these chemokines were increased in maternal blood 2-12 h (Fig.2A) their ligand-receptor interactions may occur at 2-12h, which might prevent placental inflammation by restricting immune cell recruitment and infiltration.

Notably, while CCL20 abundance in maternal blood was increased following LPS instillation at similar levels as IL-6 (Fig.2A, left), its receptor *Ccr6* was not expressed at any time point in decidua and placenta (Fig.2C, rightmost panel). Therefore, CCL20 likely does not play a role in the placental response.

As a summary, our data suggest that IL-6 from the maternal lung is a main signaling candidate for eliciting the observed immunomodulatory placental response, while placental Il6-signaling is modulated by CCL7, CCL11 and CXCL1, which may also originate from maternal liver. A necessary caveat is that we did not profile cyto/chemokine expression in all organs, and only some cyto/chemokines were measured in maternal blood^49^. Thus, cyto/chemokines could be produced by other maternal organs, and additional cyto/chemokines may have been increased in abundance in the maternal blood.

### The placenta orchestrates a time-specific response to MIA

Since the placenta reacted differently to LPS instillation than maternal lung and liver we characterized the placenta+decidua gene expression response in detail. Across all time points, 484 genes were DE (*FDR*<0.05, |log_2_FC|>0.5). The largest number of DE genes were found at 5h (313 genes; Fig.S1G). GO analysis showed distinct functional annotation enrichments for each time point, except 2h (Table S1). To investigate this further, we grouped functionally related GO terms into ‘GO themes’ (Fig.3A-F, left). For each theme, we selected a subset of genes that were linked to GO terms within the theme and among the 50 most DE at the time point where the GO theme was enriched, and visualized their LPS vs Ctrl expression change over time as heat maps (Fig.3A-F, middle).

**Figure 3:**
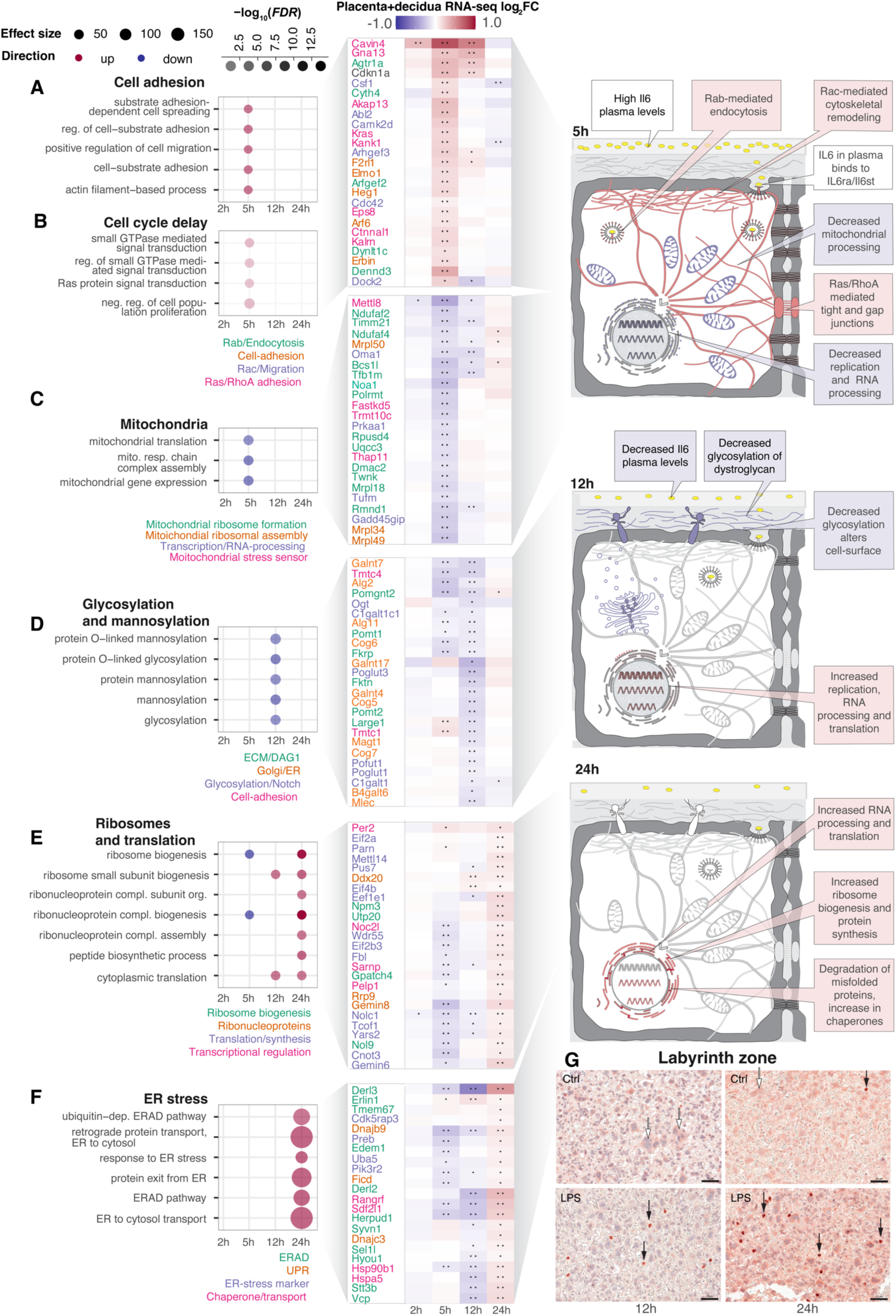
Detailed analysis of placental gene expression change. **A: Over-representation of adhesion-associated GO terms and expression change of key associated genes across time in placenta+decidua.** Left panel rows show selected adhesion-associated GO terms (cell adhesion GO theme) and their over-representation in up (red) or down-regulated genes (blue) for a given time point after instillation in placenta+decidua (X axis). Dot size and color intensity show effect size and statistical significance (-log_10_(*FDR*)). Middle panel shows placenta+decidua LPS vs. Ctrl log_2_FC expression change of top DE genes linked to one or more of these GO terms as a heat map where rows indicate genes, columns indicate time after instillation and cell color indicates LPS vs Ctrl log_2_FC. Stars indicate significant DE (LPS vs Ctrl, *FDR*<0.05). Gene names are shown to the left, colored by functional roles. Right cartoon summarizes expression changes: cell structures or organelles whose genes are up- or down-regulated as a response to LPS instillation are colored red and blue, respectively. The adhesion and cell cycle delay GO themes (panel B) have similar dynamics, therefore, genes (middle panel) are shown from both themes. Gray backgrounds link GO themes, genes and cartoon summary. **B-F: GO term over-representation and expression of linked genes for different GO themes.** Plots are organized as in A, but show different GO themes, GO terms and their associated genes. **G: IHC localization of ATF-4 expression in labyrinth zone of placenta.** Representative images of ATF-4 localization in the labyrinth zone of placenta at 12 and 24h after Ctrl or LPS exposure. Nuclei are marked by purple, ATF-4 by pink/red. White arrows indicate cytoplasmic localization of ATF-4 in trophoblast cells. Black arrows indicate nuclear localization of ATF-4 in trophoblast cells. Size bar = 50 μm.

At 5h, two GO themes were enriched in upregulated genes: i) ‘Cell adhesion’, including adhesion, spreading and migration, and actin filament-based process terms, and ii) ‘Cell cycle delay’, including small GTPase- and Ras signal transduction, and negative regulation of cell population proliferation terms (Fig.3A-B, left). Because Ras-signaling regulates cytoskeletal dynamics associated with cell adhesion and migration^50^, both themes may denote tissue-integrity functions. We therefore analyzed genes within these themes jointly. Upregulated genes were involved in cytoskeleton-mediated strengthening of tissue integrity at different levels, including Rac signaling and cell migration (e.g. *Csf1*, *Camk2d, Arhgef3, Cdc42, Dock2*), Ras/RhoA signaling and adhesion (*Cavin4, Ctnnal1, Gnai13, Kalrn, Kras*) and Rab/endosomes/endocytosis (*Agtr1a, Dynlt1c, Arfgef2, Cyth4, Dennd3*; Fig 3A,B, middle panel).

Two GO themes were enriched in downregulated genes at 5h. One encompassed aspects of ‘mitochondria’ (Fig.3C, left), including mitochondrial assembly (e.g., *Ndufaf2, Timm21, Polrmt, Dmac2, Rmnd1*), mitochondrial ribosome formation (e.g. *Mrpl34, Mrpl39, Noa1*) and sensing of mitochondrial stress (e.g., *Oma1, Prkaa1, Tufm*, *Gadd45gip1*; Fig.3C, middle). Second, genes associated with GO terms related to ‘RNA processing’ were enriched in downregulated genes (Fig.S2). In addition, ‘Ribosomes and translation’-related GO terms contained some genes that were downregulated at 5h (Fig.3E, left).

Based on the downregulation of mitochondrial production, RNA processing, ribosomes and translation genes, we hypothesized that at 5h, cell growth is reduced. We suggest that by strengthening cell-adhesion in endothelial and epithelial cell layers, and reducing changes in architecture via proliferation, the placenta restricts maternal-fetal transport of putative infectious agents^51–53^ (Fig.3A-C, right cartoon). Such adverse signals may comprise diffusion- or receptor-mediated transport of inflammatory mediators across the placental endothelium and epithelium, immune cell infiltration or translocation of microbial infectious organisms. These changes in cell architecture would largely be mediated by Ras superfamily genes, which were upregulated at 5h. This appeared to gradually decrease, as only some genes upregulated at 5h remained so at 12h (Fig.3A-C, middle).

At 12h, GO terms associated with ‘Glycosylation and mannosylation’ were enriched in downregulated genes (Fig.3D, left). One group of genes were part of alpha-dystroglycan (Dag1) signaling, either assisting in binding or glycosylation of Dag1 (*Large1, Pomgnt1-2, Pomt1-2, Fktn, Fkrp*) or as part of Notch signaling (*Pofut1, Poglut1*) which may be tuned by Dag1^54^. The dystroglycan complex links the extracellular matrix with the intracellular cytoskeleton, is regulated by glycosylation and is associated with tissue remodeling/structural changes in the placenta^55^. We speculated that reduced glycosylation of Dag1 can lead to reduced cell-matrix adhesion, ECM stiffness and tissue integrity, similar to glycosylation roles in adhesion, cell communication and infection of endothelial cells^56,57^ (Fig.3D, right). This could represent a partial reversal of the increased adhesion and tissue integrity at 5h and correlates with the decrease in maternal blood cytokine concentrations at 12h (Fig.2B). It implies reestablishment of homeostatic cell-ECM interactions and adhesion, possibly due to a decrease in sensing of infectious threats.

One GO theme pattern encompassed more than one time point: genes associated with ‘Ribosomes and translation’ terms were partially downregulated at 5h and highly upregulated at 24h (Fig.3E, S3A). At 12h, some of these genes remained downregulated while others were upregulated, suggesting a shared trend with different time trajectories. Genes following this pattern were involved in rRNA processing and ribosome biogenesis (*Nol9, Utp20, Npm3),* ribonucleoproteins (*Rrp9, Gemin8, Ddx20*), translation/protein synthesis (*Wdr55, Pus7, Eif4b, Eif2a, Eif2b3*) and transcriptional regulation (*Noc2l, Per2, Pelp1, Sarnp)* (Fig.3E).

This increase in ribosome biogenesis and protein synthesis at 12-24h was accompanied by a marked upregulation of genes involved in endoplasmic reticulum (ER) stress (Fig.3F), including genes related to ER-associated protein degradation (ERAD; *Derl2, Syvn1, Herpud1, Vcp, Erlin1*), unfolded protein response (UPR; *Dnajc3, Dnajb9, Ficd*), ER-chaperones (*Hspa5, Hsp90b1, Sdf2l1*) and ER stress marker genes (*Pik3r2, Uba5, Preb, Cdk5rap3*) (Fig.3F, middle).

The patterns of the two themes above shows the placenta’s capability of complex adaptation: the increased ribosomal, RNA processing and protein synthesis activity at 12-24h may be a compensation for the decrease of the same processes at 5h. The increase in protein biosynthesis and RNA metabolism at 12-24h may be linked to the increase in ERAD/UPR signaling and chaperone levels during the accelerated protein production in the ER lumen needed for returning to homeostasis (Fig.3E-F, right panel).

The unfolded protein response (UPR) induced during ER stress can be divided into three signaling pathways. The PERK pathway activated by the transcription factor ATF-4 initiates the mildest response, restoring ER function by attenuating non-essential protein synthesis^58^. We reasoned that a probing of ATF-4 expression in the placenta would indicate the presence of ER stress, since ATF-4 translocation to the nucleus marks the activation of the mildest ER stress pathway^59^. ATF-4 placental immunohistochemistry showed that its localisation differed between Ctrl and LPS, particularly at 12-24h in syncytiotrophoblasts of the labyrinthine zone (Fig.3G). In Ctrl, ATF-4 was exclusively cytoplasmic (Fig.3G, white arrows), except for a few nuclei per placenta at 24h (Fig.3G, black arrows). In LPS, nuclear localisation of ATF-4 (Fig.3G, black arrows) appeared in clusters of syncytiotrophoblasts throughout the labyrinthine zone, along with cytoplasmic localisation. We also stained placentas for ATF-6, a marker of the more severe ER stress pathway, but observed no obvious nuclear translocation, indicating that this more severe ER stress pathway was not activated (data not shown).

### Early placental protein phosphorylation changes reflect processes whose genes are upregulated transcriptionally

Protein phosphorylation events are essential for response to external factors^60,61^. We reasoned that part of the earliest placental response might include phosphorylation cascades and/or phosphorylation changes accompanying protein abundance changes following the observed gene expression patterns.

We measured LPS-induced phosphosite changes at 2 and 5h in placental samples where the decidua was not separated using LC-MS/MS phosphoproteomics. Few changes in phosphorylation sites occurred at 2h (216 sites in 98 proteins, *FDR*<0.05) but numbers were higher at 5h (882 sites in 578 proteins;Fig.S3A-B). GO analysis showed that changes primarily occurred in proteins mirroring the functional roles of genes upregulated at 5h (Fig.3, S3), in particular proteins associated with DNA metabolism, transcriptional regulation, cell growth and RNA/protein processing or metabolism (Fig.4A). We also observed over-representation of damage response terms, in particular viral processes and DNA damage. The two latter themes may be reflections of the first, i.e. phosphorylation changes in DNA damage proteins may reflect regulation of growth processes.

**Figure 4:**
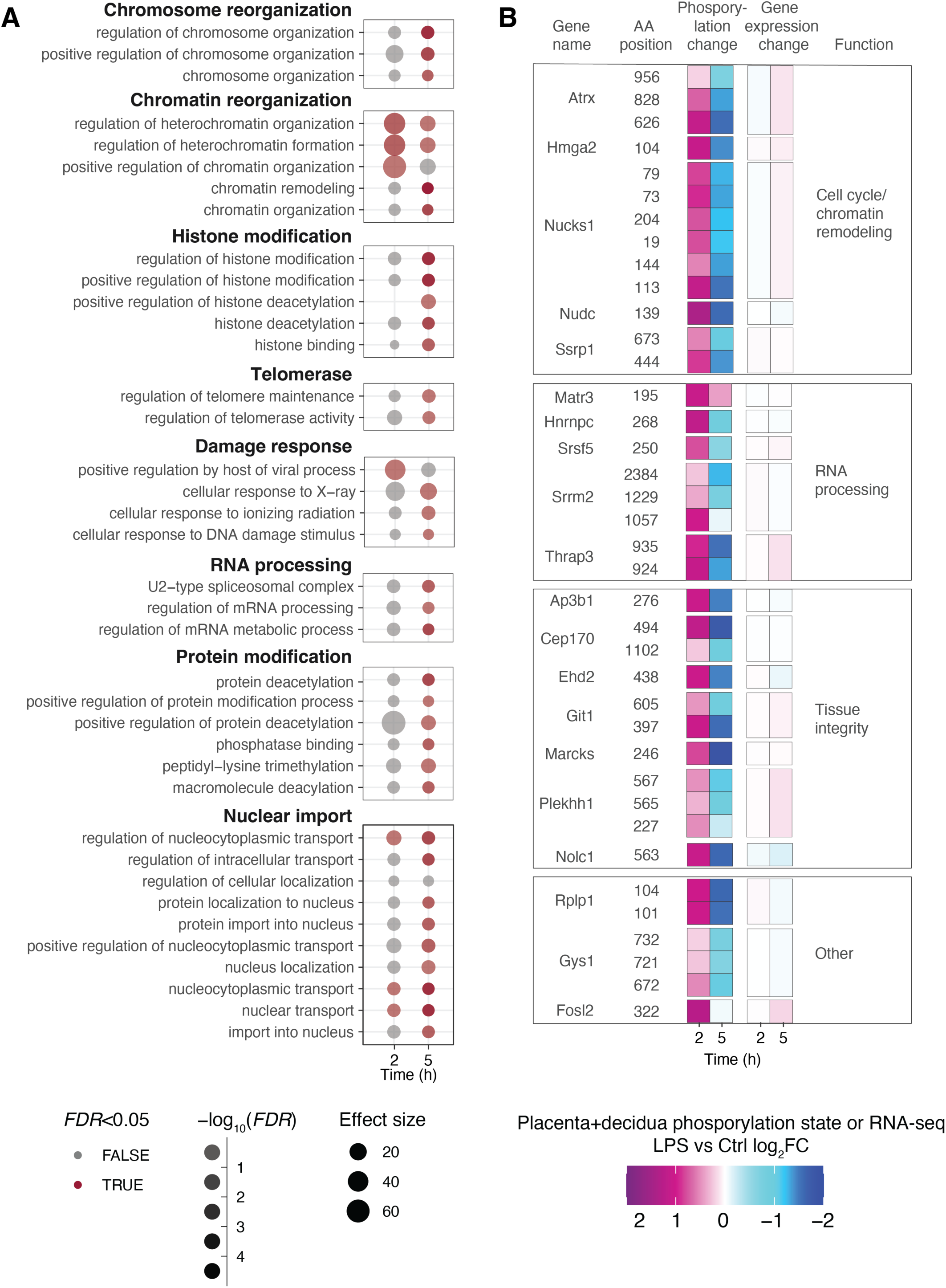
Phospoproteomics analysis of early placental response. **A: Gene ontology analysis based on genes with changing phosphosites in placenta.** Rows show GO terms, organized in functional themes as indicated on top of boxes. Columns indicate time after instillation (h). Dot opacity indicates significance (red color indicates *FDR* <0.05), dot size indicates effect size. Analysis is based on proteins having one or more significant phosphosite changes, regardless of direction of change. **B: Proteins and phosphosites with LPS-induced phosphorylation increase at 2h and decrease at 5h.** Rows in the left two columns of the heat map show phosphosites for a given protein at 2 and 5h, as indicated by two first columns. Colors show average LPS vs Ctrl log_2_FC, where positive values represent a gain of phosphorylation in LPS vs Ctrl. The two columns to the right show change in expression level LPS vs Ctrl log_2_FC for the corresponding gene at 2 and 5h. The last column shows the gene function category. Displayed phosphosites have a significant (*FDR*<0.05) change in phosphorylation state at 2 or 5h, a LPS vs Ctrl log_2_FC >0 at 2h, and higher LPS vs Ctrl log_2_FC at 2 than 5h.

Interestingly, several phosphoproteins had increased phosphorylation at 2h but decreased phosphorylation at 5h, comparing LPS and Ctrl (Fig.4B, left). Notably, many of these proteins have functional roles in tissue integrity, RNA processing and chromatin remodeling (Fig.4B, left); themes that were also enriched in upregulated genes at 5h on RNA level (Fig.3, S3). Despite this similarity on GO and hence functional level, proteins with these phosphorylation patterns were not DE at mRNA level (Fig.4B, right). This suggests the existence of a protein phosphorylation wave occurring earlier than changes at RNA expression level and affecting other proteins than those being transcriptionally upregulated. Consistently, phosphorylation of several proteins with this pattern (e.g. Atrx, Hmga2, Nudc) is cell cycle dependent^62–64^, and, as noted above, genes associated with cell cycle delay were upregulated at 5h.

In summary, we did not detect substantial phosphorylation changes indicative of early response pathway phosphorylation cascades. Instead, changes occurred mainly in proteins with functions similar to those of the differentially expressed genes. Thus, placental+decidual response likely occurs at both protein and RNA level.

### Maternal immune activation leads to metabolic adaptation in fetal liver

In fetal liver 755 genes were DE (*FDR*<0.05, |log_2_FC|>0.5) at ≥1 time points. We observed several similarities to the placental response. First, most genes were DE only at one time point, with most DE genes at 5h (473, Fig.S1G). Second, GO analysis showed distinctive functional enrichments for each time point. As above, we grouped related GO terms into GO themes (Fig.5A-E, left), and visualized expression changes of top DE GO-linked genes (Fig.5A-E, middle). Notably, we observed no enrichment of immune response terms.

**Figure 5:**
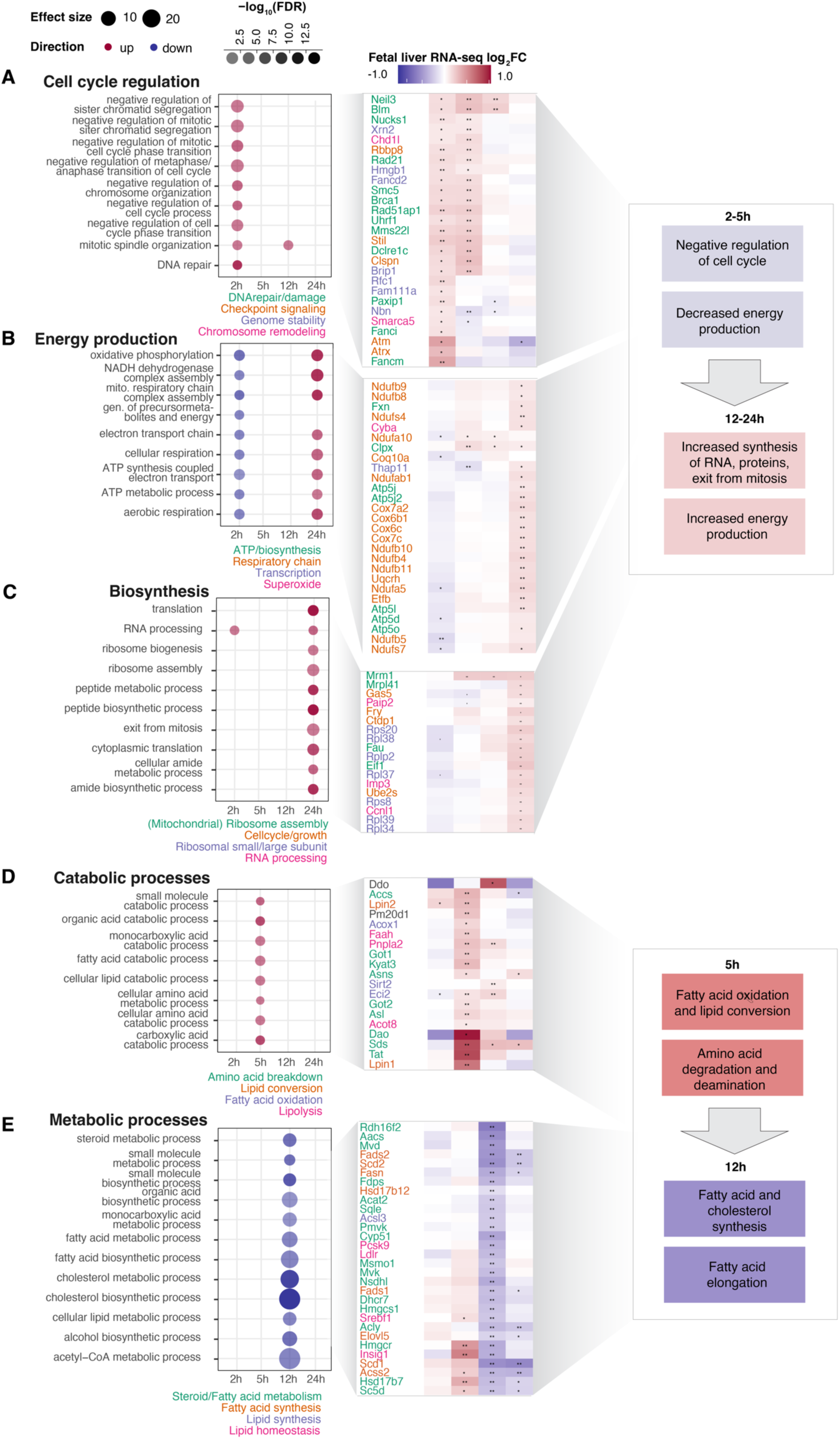
Detailed analysis of fetal liver gene expression change. The figure is organized as Fig.3A-F but based on fetal liver RNA-seq data. Boxes to the right summarizes expression changes early and late in the time course.

One larger pattern was reminiscent of the placental response: early down-regulation of ‘energy production’ and ‘cell cycle regulation’, followed by upregulation of ‘energy production’ and ‘biosynthesis’ themes at 24h. Specifically, the fetal liver upregulated genes associated with negative regulation of cell cycle at 2h (Fig.5A, left), including genes with roles in DNA repair (e.g. *Fancm, Neil3, Blm, Rad51ap1, Paxip1*) and genome or chromosome stability (e.g. *Brip1, Fancd2, Hmgb1, Nbn, Fam111a)*. Some of these genes were also upregulated at 5h (Fig.5A, middle). Genes annotated with GO terms related to ‘energy production’, including cellular and aerobic respiration, oxidative phosphorylation and mitochondrial processes (Fig.5B, left) were subtly downregulated at 2h, at similar expression levels as Ctrl at 5-12h, and substantially upregulated at 24h (Fig.5B, middle). These genes included *Nduf*-, *Cox-* and *Atp*-family genes involved in the respiratory chain. Related to this, at 24h, biosynthesis (including translation, RNA processing, ribosome biogenesis/assembly) and protein/amide synthesis GO terms were upregulated, e.g. large and small ribosomal protein subunit genes (*Rpl-* and *Rps-*genes) and cell cycle progression genes (e.g. *Ube2s, Ctdp1*;Fig.5C).

Overall, these observations imply that the fetal liver responds to LPS by depleting/inhibiting the translational machinery, cell division, and energy production capacity at 2h, followed by a gradual increase of these processes until at 24h where the cell cycle machinery is restored, and energy and RNA/protein/ribosome biosynthesis gene expression is highly increased (Fig.5A-C, right). We hypothesize that at 24h, the fetal liver reestablished high capacity for biosynthesis, along with needed energy production systems, to compensate for the decreases in cell cycle progression and biosynthesis at earlier time points. This pattern resembles that of the placenta, although placental upregulated genes covered a broad range of functions related to protein synthesis and ribosome biogenesis, while fetal liver upregulation related almost exclusively to ribosome biogenesis and mitochondrial energy production.

One gene expression pattern unique for the fetal liver was upregulation of ‘catabolic processes’ at 5h and downregulation of ‘metabolic processes’ at 12h. Specifically, genes associated with amino acid breakdown (including *Tat, Sds, Got1, Dao* and *Asns*,) lipid conversion (including *Lpin1-2*), fatty acid oxidation (including *Sirt2, Eci2* and *Acox1*), and lipolysis (including *Faah, Pnpla2* and *Acot8*) were upregulated at 5h. Many of these genes remained slightly downregulated at 12h (Fig.5D, right). At 12h, genes associated with metabolism-related GO terms were downregulated, including genes with key roles in steroid/fatty acid metabolism (e.g. *Fads2*, *Hmgcs1*, *Hmgcr*, *Acly*, *Insig1*, *Acat2*), lipid synthesis (e.g. *Slc27a3*, *Gpat4*, *Ppard*, *Acsl3*, *Fads1*) and regulation of fatty acid elongation (e.g. *Hacd3*, *Hsd17b12* and *Elovl* family genes). Many of these genes use acetyl-/Acyl-CoA for synthesis of lipids. Acetyl-CoA is synthesized in mitochondria during cellular respiration and cellular respiration genes were downregulated at 5h (Fig.5C). Acetyl-CoA availability is determined by the metabolic status of the cell: during fasting, the more Acetyl-CoA is needed for ATP generation in mitochondria, the less is available for cytosolic lipid synthesis^65^.

Therefore, we hypothesized that the observed pattern reflected a substantial change in metabolic state due to lack of nutrients. The fetus does not synthesize glucose but depends on maternal supply^66^. Similarly, maternally derived long-chain polyunsaturated fatty acids (LC-PUFA) are essential for fetal brain development^66^. We suggest that the fetus’ availability of glucose and LC-PUFA was briefly decreased, based on net weight loss in LPS-instilled dams vs. Ctrl (Fig.S1D), and that the fetal liver gene expression response reflects a shift to alternative pathways than glucose oxidation for energy generation. Possibly related to these metabolic adaptations, genes associated with glycosylation-related and development GO-terms were downregulated at 5h (Supplementary Note S2-3, Fig.S4).

### Maternal LPS exposure increases levels of essential fatty acid carriers in fetal liver

We reasoned that lipid metabolism gene expression changes would be reflected in lipid abundance. LC/MS lipidomics analysis of fetal liver samples from all timepoints detected 1047 lipid species, where 189 differed significantly between LPS and Ctrl (*FDR*<0.05, ebayes test^67^, Benjamini-Hochberg correction) at ≥1 timepoints. A heatmap of the 50 most significantly changing lipids revealed two outstanding patterns (Fig.6A). First, at 12 and 24h, levels of a large set of lipids containing docosahexaenoic acid (DHA; 22:6 ω-3) were increased in LPS vs Ctrl. DHA-containing lipids primarily comprised triglyceride (TG)–species containing one to three DHA side-chains. Levels of other lipids with one or more DHA side-chains were also increased, including diglycerides (DG), phosphatidylglycerol (PG), phosphatidylcholine (PC), phosphatidylethanolamines (PE) and fatty acyl esters of hydroxy fatty acid (FAHFA) (Fig.6A). Second, at 5 and 12h, levels of diverse lipids decreased in LPS vs Ctrl. None of these contained DHA and most were phospholipid-membrane species.

**Fig 6:**
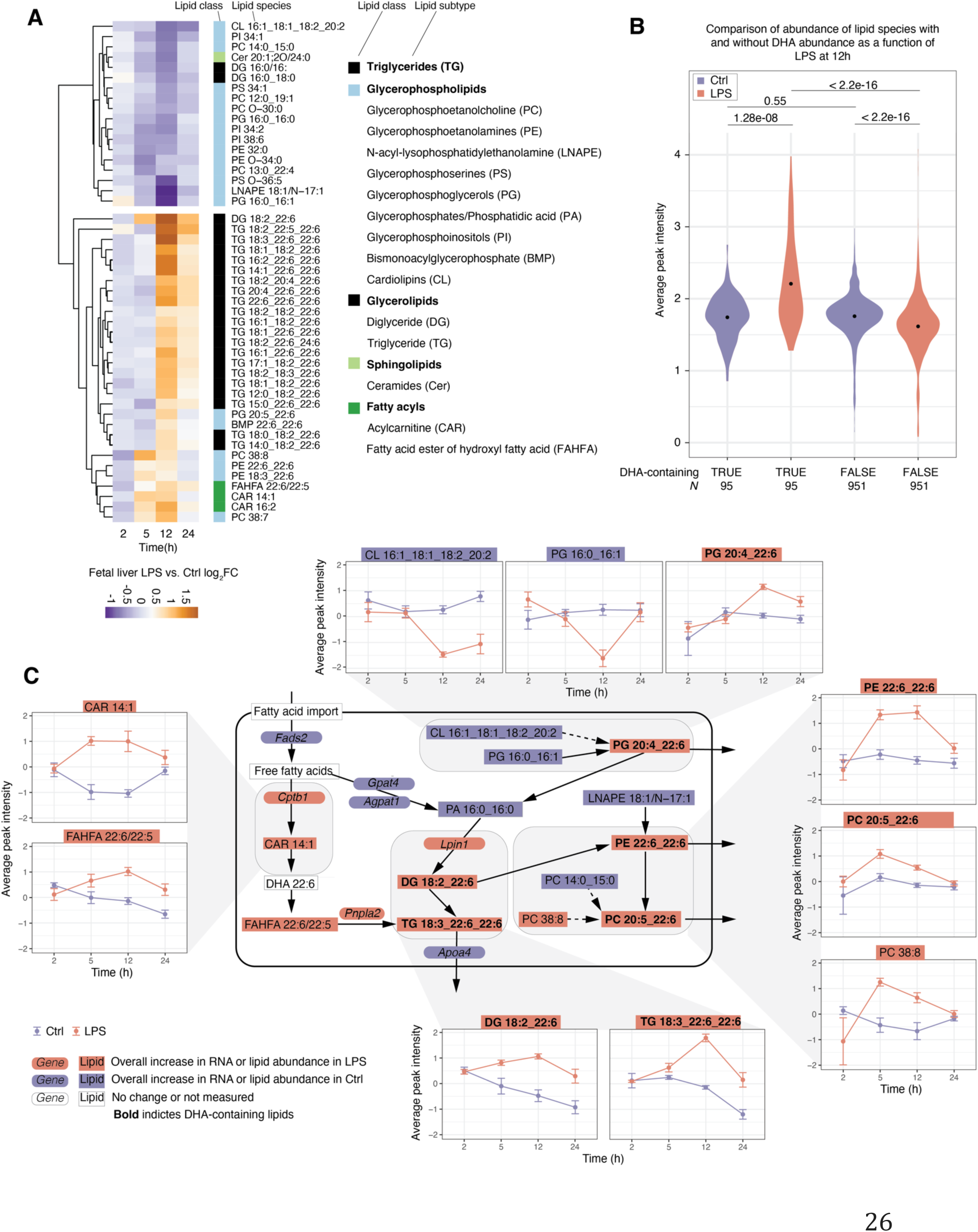
Lipidomics analysis of fetal liver. **A: LPS vs Ctrl log_2_FC of lipid abundance at each time point.** Heat map is based on the top 50 most significantly LPS vs Ctrl changing lipid species (sorted by lowest *FDR* at any time point) clustered by euclidean distance. Lipids are annotated by class (right). **B: Lipid abundance as a function of whether lipids contain DHA and LPS/Ctrl treatment.** The Y axis shows average peak intensity. Numbers on top are *P*-values from Whitney-Mann two-sided tests. Number of lipid species in each group is indicated at the bottom. Dots within density plots indicate distribution means. **C: Network of selected lipids and lipid–processing enzymes.** Arrows show conversions of lipids or lipid classes (boxes: bold typeface indicate DHA-containing species). Selected enzymes for conversion are shown (rounded boxes). Boxes are colored by whether the lipid or enzyme is up-down or unchanged over time following LPS instillation. Examples of temporal regulation on lipid species levels are shown as callouts: in these, Y axis show average abundance (peak intensity), X axis show time after instillation. Error bars show SEM. Arrows ending outside the cell boundary denote putative export of lipids into the bloodstream by lipoproteins.

More generally, we found significantly increased abundance of DHA- vs non-DHA-containing lipids in LPS but not Ctrl at 12h (*P*=1.281e-08 and 0.5489, respectively, two-sided Wilcox test). Furthermore, levels of non-DHA-containing lipids were significantly lower in LPS than Ctrl at 12h (*P*<2.2e-16, two-sided Wilcox test) (Fig.6B).

Although untargeted lipidomics cannot detect free DHA, our observations suggest a global increase in lipids with incorporated DHA in response to LPS instillation. While the maternal diet is the main factor determining fetal brain DHA-accretion, fetal liver biosynthesis and storage of DHA is also important^68,69^, as it can be secreted to fetal plasma within lipoproteins.

We drew a simplified network of lipids with high abundance changes and associated lipid processing enzymes, and plotted lipid abundance (Fig.6C, callouts for selected lipids, Fig.S5A) and gene expression (Fig.S5B) change over time. This enabled four observations:

First, phospholipids that could be converted into DHA-containing phospholipids were typically downregulated at later time points with a reciprocal increase of downstream DHA-containing phospholipids (e.g. PG 14:0_15:0→PC 22:6_22:6/PC 20:5_22:6 and PG 16:0_16:1→PG 20:4_22:6, Fig.6C, top).

Second, enzymes for elongation and conversion of fatty acids into phosphatidic acid (PA) (PA 16:0_16:0), a key intermediate in lipid-synthesis, including *Fads2, Gpat4 and Agpat1,* were downregulated (Fig.S5B).

Third, nearly all lipids downstream of PA containing DHA were upregulated, as were many of the enzymes that catalyze lipid conversions in this part of the pathway (e.g *Cptb1, Pnpla2, Lpin1*; Fig.6C, left and bottom, S6B).

Fourth, DHA-containing lipids stored in lipid droplets or poised for exported into the bloodstream via lipoproteins were highly upregulated at 5-24h, commonly most pronounced at 12h (Fig.6C, right).

Thus, it is possible that the source of increased DHA-containing lipids after LPS installation was stored fetal liver lipids rather than imported. The decrease in several phospholipids which can be broken down as sources for DHA-containing species supports this notion (Fig.6A). In addition, as discussed above, many enzymes that use Acetyl-CoA to generate energy or for *de novo* fatty acid synthesis were downregulated at 12h (Fig. S6C).

As a summary, we suggest that the fetal liver upregulated production of DHA-containing lipids from existing lipids because of decreased maternal lipid- and glucose influx. Notably, fetal liver storage of DHA functions as a deposit for other fetal organs, and fetal liver can export DHA to the bloodstream^69^, particularly during the first hours of postnatal neuro-development^66,69,70^, where brain-accretion of DHA is highest. We could not carry out lipidomics on fetal blood to establish whether the increase in DHA-containing lipids primarily leads to storage or export.

## DISCUSSION

Here, we analyzed responses to LPS-induced MIA in maternal lungs, across maternal, placental and fetal tissues and time, based on transcriptomics, phosphoproteomics and lipidomics, complemented by targeted protein abundance assays of maternal blood and placental imaging.

While LPS induced a clear immune response in maternal lung and liver, through activation of TLR4-pathway genes (Fig.1B-C) and increased levels of cyto- and chemokines in maternal blood (Fig.2A), this did not extend to placenta and fetal liver which displayed highly different, and time-specific responses.

Specifically, placenta increased expression of tissue integrity genes while DNA/RNA processing, biosynthesis and energy production gene expression decreased at 5-12h, followed by compensatory increases in protein synthesis and ribosome biogenesis gene expression and, at 24h, mild ER-stress. The placenta did not mount an innate immune response, even though it has the ability in direct response to LPS^34,35^. This could be due to lack of direct LPS exposure. However, despite that maternal liver and placenta are expected to be exposed to similar LPS levels, maternal liver did display a pronounced innate immune response, including upregulation of TLR4-pathway genes. Notably, the few upregulated placental TLR4/IL-6-pathway genes, e.g. *Nfkbiz* and *Socs3,* modulate inflammation rather than activate innate immune response. Related to this, our data indicate that IL-6 originating from maternal lungs is the most likely candidate to elicit this response, although interestingly, the placenta did not increase *Il6* production itself. In late gestation, the placenta is almost impermeable to IL-6^71^ and IL-6 signaling from mother to fetus at late gestation therefore depends on placental IL-6 production^13^. As discussed above, *Socs3* was upregulated in the placenta and is an inhibitor of IL6 production via the Jak/STAT-pathway. It therefore might have prevented the induction of inflammatory cascades. This observed capability of selective ‘immune inertia’ may be a feature of placental immune adaptation^72^ and immune tolerance^73,74^, possibly linked to the pulmonary rather than intravenous LPS exposure.

The fetal liver, somewhat similarly, first down-regulated cell cycle signaling and growth genes, but increased growth and protein synthesis gene expression at 24h. At 5-12h, levels of DHA-containing TGs and PEs were increased, likely by conversion of existing lipids. DHA deposited in fetal liver is known to buffer decreased availability from the mother^69^ and suggested to prepare offspring for parturition and the excessive DHA-demands for neonatal brain development.

In summary, the placenta selectively avoided an innate inflammatory immune response and instead initiated a time-specific adaptation program that appeared to not transfer innate immune response signaling cues to the fetus, based on the lack of innate immune response in the fetal liver. On the other hand, the fetus experienced a temporary drop in nutrients from the mother leading to conversion of existing lipids to DHA-containing TGs and PEs. This adaptive response was remarkably transient: after 24h, placenta and fetal liver gene expression in LPS and Ctrl were largely similar.

Our study has important limitations. First, it only assesses short-term MIA effects. Long-term effects could have manifested in the offspring of LPS-treated mothers, since maternal LPS treatment, albeit via intraperitoneal or intravenous routes, can have severe long term effects in offspring organs^9,10^. Second, our study is descriptive, and although our interpretations in many cases are supported by multiple experiments, they are not causally proven.

Nevertheless, to our knowledge, the overall temporal maternal and fetal response, adaptation and return to homeostasis after MIA has never been described previously using omics methods. Our study sheds new light on this process and the adaptive capabilities of the placenta, in particular during distal MIA.

## MATERIALS AND METHODS

### Animals and ethics

80 Nulliparous C57BL/6JRj mice (Janvier, Saint Berthevin Cedex, France) from the same barrier room were time-mated and pregnancy confirmed by the presence of vaginal plug the morning after mating (designated gestation day (GD) 0). Dams arrived at the institute at GD 11 or 12 and acclimated for 6-7 days prior to exposure. Mice were pair housed in clear 1290D euro standard polypropylene cages with Aspen bedding (Tapvei, Estonia), enrichment (mouse house 80-ACRE011, Techniplast, Italy; small aspen blocks, Tapvei, Estonia) and nesting material (Enviro Dri, Lillico, Biotechnology, UK), under controlled conditions (temperature 21-22°C; humidity 55+/-10%; ventilation 15-20 air changes/hour; 12 hour light-dark cycle with lights on at 06.00 a.m.) and access to food (Altromin 1314 for breeding, Brogaaarden, Denmark) and tap water ad libitum. Animals were weighed on the day of arrival and the day prior to exposure to confirm pregnancy. All procedures followed the guidelines for care and handling of laboratory animals established by the EC Directive 86/609/EEC and Danish regulation (Danish Ministry of Justice, Experimental Animal Inspectorate, permit 2015–15–0201-00569). The local animal welfare committee approved the specific protocol prior to the study.

### Exposure

Lipopolysaccharide (LPS; E. Coli serotype 00:55 B5 LPS (Sigma Lot nr. 025M4040V)) was diluted to the final concentration (0.02 µg/µl) in double distilled pyrogen-free water (Chem-Lab, Zedelgem, Belgium). In the morning of GD 17, the pregnant mice were semi-randomized into control and LPS treatment groups (denoted Ctrl and LPS, respectively), evenly distributing weights among the groups. Out of 80 mice in total, 74 were pregnant. Animals were placed in a whole-body inhalation chamber with an attached anaesthetic vaporizer (Penlon Sigma Delta, Abingdon, UK), delivering 3-4% isoflurane in filtered air, and were intratracheally instilled with 50 µl of vehicle (Ctrl) or 1 μg LPS in 50 µl vehicle, followed by 200μl of air. Vehicle and LPS were administered through a 0.58 mm polyethylene tube (Ref: 427411, Becton Dickinson, Brøndby, Denmark) attached to a plastic syringe. The procedure has been shown not to affect gestation, offspring viability nor growth^75^. After instillation, animals were returned to their cage, briefly placed on heating pads and checked upon regularly until euthanization.

### Isolation of pups and maternal/fetal organs

At 2, 5, 12 and 24h, dams were terminally anesthetized by subcutaneous injection of 0.2 ml of Zoletil mixture (tiletamin/zolazepam, xylazin og fentanyl) and killed by exanguination by withdrawal of heart blood into Eppendorf tubes containing 36 ml K_2_EDTA (*N*=7-9 per exposure/time point). The uterus was excised and opened. Fetuses were excised from their embryonic sac, their viability confirmed, killed by decapitation, sexed by visual inspection, and their position in the uterus noted. From each litter, the first female fetus encountered in the right uterine horn, counting from the cervix, was selected and saved for analyses. The placenta was dissected into chorion (chorionic plate, labyrinth and junctional zones) and decidua by blunt/stump dissection under stereomicroscope (Wild Heerbrugg, Switzerland)^76^. From dams, the liver and right lung were dissected. Dissected organs were snap frozen in liquid nitrogen and kept at −80°C. Fetal livers were excised later. Maternal blood was centrifuged at 2000 x g at 4°C for 5 min and plasma was stored in aliquots at −80°C until analysis. A maximum of one female fetus from each dam were used for any one outcome, except for lipidomics, where two female fetuses were used for some of the groups/time points.

### Fetal sex genotyping

Genotyping of pup sex was performed on DNA extracted from the fetal tail, by PCR using primers for *Ddx3Y* (denoting the Y chromosome in males: forward 5’-GGG TCT GTG ATA AGG ACA GTT CA-3’, reverse 5’-CAC GAC CAC CAA TAC CAT CAT AG-3’) and *Rpl13a* (denoting the X chromosome present both in males and females, forward 5’-AGC CTA CCA GAA AGT TTG CTT AC-3’, reverse 5’-GCT TCT TCT TCC GAT AGT GCA TC-3’), purchased from TAG Copenhagen A/S. Following PCR, the reaction was separated on a 1.5% agarose gel. A band for Ddx3Y of 908 bp, connoted XY whereas lack of this band but presence of Rpl13a of 129 bp, connoted XX.

### RNA extraction and library construction

Total RNA was isolated from frozen maternal lung and liver, chorion, decidua and fetal liver. Briefly, 20-50 mg of tissue was homogenized with a T 10 basic ULTRA-TURRAX® blender (IKA, Staufen, Germany) in 700 ul lysis buffer with 7 ul mercapto-ethanol. RNA extraction was carried out utilizing magnetic beads technology, on a chemagic Prepito® (Perkin Elmer, Waltham, Massachusetts), as recommended by the manufacturer. Concentration and purity were measured on a NanoDrop1000 spectrophotometer, with all samples showing an A260/280 ratio between 1.9 and 2.1. RNA integrity was analyzed by 2100 Bioanalyzer (Agilent Technologies) with Agilent RNA 6000 Pico Kit (Agilent Technologies) as recommended by the manufacturer. All samples used for RNAseq displayed RNA integrity number (RIN) above 7. cDNA library construction and paired end sequencing was carried out by Novogene (China). To exclude ribosomal RNA, polyA selection was done.

### RNA-seq analysis

RNA-seq analysis from quality control to DE analysis was done using a Snakemake^77^ pipeline using Conda (https://conda.io). Code and parameters are available at https://github.com/drdariarago/INFLAMMATION_TRANSFER. Sequencing produced a total of 740 libraries (370 paired end), with a median read depth of 24 million reads (mean 25 million reads, min 19.9 and max 46.6), all of which passed initial QC controls performed by multiqc^78^. We detected base-pair overrepresentation in the first 11 bp of each read, as expected due to biases in random primers, and removed them using the seqtk (https://github.com/lh3/seqtk). We used Salmon^79^ to map reads to gencode mouse transcriptome version M23^80^, which is an annotation of the genome assembly version GRCm38. For mapping, we created an index using *k*-mer size 31 bp and supplying the genome in order to create decoy *k*-mers to account for biases due to un-annotated transcribed genomic regions (as described in Salmon documentation). We mapped libraries to the resulting index using Salmon quant using the following options: gcBias, seqBias, validateMappings, numBootstraps=10 and minScoreFraction=0.8. Finally, we annotated the resulting quant files using the R library tximeta^81^ and removed all features that lacked annotation. We performed all initial exploratory analyses and plots using TPM-normalized data, and used the raw count data for the differential expression analysis. Before fitting statistical models, we normalized the count data from Salmon using TMM, retained only transcripts with >10 reads in at least 70% of the samples of the same condition^67^. We detected differentially expressed genes using generalized linear models in limma after variance stabilization using voom^67^. We converted all *P*-values to *Q*-values using FDR correction using FDRtool^82^. Due to the large differences in expression between most tissues (Fig.S1E) we fitted independent models for each tissue, with the exception of placental samples (see below). For maternal lung, maternal liver and fetal liver we used the model formula *E = 0 + timepoint + timepoint:LPS* which estimates each gene’s average expression for control samples in each timepoint, and then estimates the difference in expression between control and treatment samples from the same time point. Since we found that placenta (encompassing chorionic plate, labyrinth and junctional zones) and decidua were very similar in their overall expression response (Fig.S1J) we fitted a single model for both, with the formula *E = 0 + timepoint + maternal + timepoint:LPS + timepoint:LPS:maternal.* This model calculates the average gene expression at each timepoint for control samples of both placenta and decidua, and then tests for (1) differences in the expression of decidua and placenta within control samples, (2) shared responses to maternal inflammation and (3) differences between the responses to maternal inflammation of both. Although several genes showed differences between placenta and decidua (contrast 1), we found that only 8 genes showed significant differences (*FDR* < 0.05) in their response to maternal inflammation between the tissues with an absolute log_2_FC >1 (contrast 3), and 7 of those were unannotated transcripts.

### Gene set enrichment analysis

We performed gene set enrichment using leading-edge analysis using gprofiler2^83^. For each tissue, we selected all genes with baseline normalized fold expression >0 and ordered them by *Q*-values, signed according to up or downregulation (decreasing to test for upregulation, increasing to test for downregulation). We used the list of all genes expressed in each tissue above log_2_ normalized counts of zero in at least one time point as the background set for all enrichment tests in that tissue. We used the gSCS method for *P*-value correction with a threshold of 0.05 and testing only for over-representation of GO terms. In order to summarize the results for figures, we retained only significant Biological Process terms with more than 10 and less than 1000 terms. Due to the vastly different amount of differential expression between tissues, we used different methods to summarize the results. From the maternal lung and liver GO terms, we curated lists of highly significant (*FDR* <0.01 at any time point) GO terms that were both interesting and representative, which were used for Figure S1I-J. Similarly, for the placenta and fetal liver GO terms, we manually curated a list of interesting and representative GO terms (FDR <0.01 and effect size > 2, at any time point). From these, we extracted the complete list of genes annotated with the respective GO term(s), and only retained those genes that were differentially expressed (*FDR* <0.01) at the time point where the GO-term was significantly enriched, and from these retained the 50 most significant genes sorted by FDR.

### Cytokine abundance measurements in maternal blood

A total of 31 chemokines were measured in maternal plasma using a magnetic bead-based kit (Bio-Plex Pro Mouse Chemokine 31-Plex). The Luminex xMAP multiplexing technology and the Bio-Plex® 200 platform (Bio-Rad Laboratories, USA) were used for analysis of the plasma samples. Plasma was diluted 1:4 and the protocol carried out according to the manufacturer’s description. The standard curve was run in duplets, the samples in singlets. After initial analysis of plasma protein concentrations, 10 out of the 31 chemokines were chosen for further analysis.

### Ligand-receptor analysis

We used the CellPhoneDB database^43^ annotations to link annotated secreted proteins/peptides with cognate annotated receptors. The linkage allowed for many-to-one and many-to-many matches (e.g. several ligands bound to one receptor, or vice versa). Since the cellphone database is human-based, we translated the human gene names to their orthologous mouse counterparts using Ensembl^84^ annotation.

### Immunohistochemistry

For immunohistochemical staining, placental tissue from female pups was fixed in paraformaldehyde (PFA 4%, overnight) before embedment in paraffin. Sections were cut to a thickness of 7 μm with a Leica HistoCore AUTOCUT Rotary Microtome (Leica Biosystems, Wetzlar, Germany) and Leica RM2245. The remaining experimental protocol was carried out on a Bond RXm (Leica Biosystems, Wetzlar, Germany) staining robot. At this point, sections were dewaxed with Bond Dewax solution and boiled for target retrieval in BOND Epitope Retrieval Solution 2 for 20 min. Between each step, sections were washed with Bond Wash Solution. Endogenous peroxidase activity was blocked with 3% H_2_O_2_ for 5 min, followed by blocking by incubation in 10% donkey serum (Candor Bioscience, Wangen, Germany) for 10 min. The primary antibody for ATF-4 (dilution 1:100, Abcam/ab31390, Anti-rabbit) was incubated 1h at ambient temperature. Sections were coated with a biotinylated secondary antibody (dilution 1:500, donkey anti-rabbit, reference: 711-065-152, Jackson ImmunoResearch) for 30 min and incubated with ABC solution (Vectastain PK-6100; Vector Laboratories, Linaris, Dossenheim, Germany) at room temperature for 30 min. Finally, the AEC kit (SK-4205; Vector Laboratories) was applied, and sections were counterstained with hematoxylin for 15 seconds and embedded in Vectamount AQ Aqueous Mounting Medium (reference: H-5501, Vector Laboratories). Slides were scanned using a PANORAMIC SCAN (3DHISTECH Ltd., Budapest, Hungary, and analyzed with the CaseViewer software (Version 2.4, 3DHISTECT Ltd.).

### Proteomics sample preparation, TMT labeling and chromatography

Female placenta samples from 5h (16 samples in total) were subjected to lysis with 5% SDS in water at room temperature, sonication and boiling for 10 minutes at 95°C Protein concentration was measured, and 200 µg of sample were processed to reduction, alkylation, and digestion with LysC and Trypsin enzymes by the Protein Aggregation Capture (PAC) method as in^85^ using MagReSyn® HILIC microparticles (ReSyn Biosciences Ltd). 100 µg of tryptic peptides from each sample were used for the TMTpro reagent (ThermoFisher Scientific) labeling procedure according to the manufacturer’s protocol. The TMT-labeled peptides were pooled together, lyophilized, and applied to a phosphopeptide enrichment protocol^86^ by immobilized metal-ion affinity chromatography (IMAC) with MagReSyn® Ti-IMAC magnetic microparticles (ReSyn Biosciences Ltd). The eluted from IMAC peptides were subjected to the High pH fractionation as in^87^ resulting into 14 HpH fractions that were dried out in a vacuum centrifuge and resuspended in 0.1% trifluoroacetic acid (TFA) for subsequent LC-MS/MS.

### Phospho-proteomics LC-MS/MS, raw data processing and analysis

The resulted samples were infused into the home-made fused silica column (inner diameter of 75 µm) packed with C18 resin (1.9 µm beads, Reprosil, Dr. Maisch) with an EASY-nLC 1000 ultra-high-pressure system (Thermo Fisher Scientific) for reverse phase chromatography. A high-field asymmetric waveform ion mobility spectrometry (FAIMS Pro) device (Thermo Fisher Scientific) was placed between a nanoelectrospray ion source and an Orbitrap Exploris 480 mass spectrometer (Thermo Fisher Scientific). The FAIMS was operated in a standard resolution mode with CVs −50V and −70V that were applied to all scans of the entire MS run. The Orbitrap Exploris 480 MS was used in positive-ion mode with a capillary temperature of 275°C acquiring MS data in a data-dependent mode (DDA) based on cycle time with master scans equal to 1.5 second. Method duration was 180 minutes with a normalized AGC target value 300% at full MS scan. Resolution was set to 120000 with scan range 400-1400 m/z and maximum injection time (IT) 50 ms. The Normalized Collision Energy (NCE) by HCD was 32%. For the MS/MS scan resolution was set to 45000, maximum IT to 120 ms, isolation window with 0.7 m/z, normalized AGC target was 200%. The dynamic exclusion window was set to 60 s. The resulting 14 raw files were processed to MzXML files using FAIMS MzXML generator (https://github.com/coongroup/FAIMS-MzXML-Generator) to search 28 MzXML files with MaxQuant (version 1.6.7.0) applying TMTPro correction factors for TMT channels quantitation by the software. The search was done against a target/decoy database (Mus Musculus, SwissProt from September 2019 with 17013 entries) with *FDR* < 0.01 with the following parameters: main search peptides tolerance was 10 ppm, fragment mass tolerance was 20 ppm. An enzyme for protein digestion was specified as trypsin with allowing twp missed cleavages. Cysteine’s carbamidomethylation was specified as fixed modification and protein N-terminal acetylation, oxidation of methionine and phosphorylation on STY residues were set as variable modifications. Results from the MaxQuant search “Phospho (STY) Sites” table were used for identified phosphosites quantitation analysis using the edgeR package^88^ using TMM-based normalization and differential abundance analysis using *FDR*<0.05 as a significance threshold. Gene set enrichment analysis of proteins with changing phosphorylation states was made in the same way as RNA-seq data, with the following changes: as input, we selected all proteins with one or more changing phosphosites (*FDR*<0.05, as defined above), and for each protein, we only retained the lowest *FDR* value if several sites satisfied this criterion. We then ordered these proteins based on *FDR* and used this list as input for gprofiler2. As background, we downloaded a list of all *M musculus* proteins with one or more experimentally validated phosphosites from the EPSD database version 1.0^89^, and then intersected this with RNA expression data from the same tissue, only keeping genes/proteins which were detected by RNA-seq and having one or more phosphosites.

### LC/MS lipid profiling, data processing and analysis

Lipids were extracted from fetal liver samples (20 mg) using Folch extraction^90^ with 8-12 replicates from each experimental group at each time point. Prior to tissue lysis, Splash mix (Merck) was added to the extraction solvent and tissue samples were lysed by beat beating in a FastPrep-24 homogenizer. After centrifugation and phase separation, the apolar and polar phases were transferred to separate tubes, and the apolar phase dried under N_2_. Samples were resuspended in 30 µl methanol/chloroform (1:1) and centrifuged (5 min/16,000×g/22°C) before transferring to HPLC vials. A quality control sample was constructed by pooling 3 µl of each sample. Samples (0.5 µl) were injected using an Vanquish Horizon UPLC (Thermo Fisher Scientific) equipped with a Waters ACQUITY Premier CSH (2.1 x 100mm, 1.7 µM) column operated at 55°C. The analytes were eluted using a flow rate of 400 μL/min and the following composition of eluent A (Acetonitrile/water (60:40), 10 mM ammonium formate, 0.1% formic acid) and eluent B (Isopropanol/acetonitrile (90:10), 10 mM ammonium formate, 0.1% formic acid): 40% B from 0 to 0.5 min, 40–43% B from 0.5 to 0.7 min, 43-65% B from 0.7 to 0.8 min, 65-70% B from 0.8 to 2.3 min, 70-99% B from 2.3 to 6 min, 99% B from 6-6.8 min, 99-40% B from 6.8-7 min before equilibration for 3 min with the initial conditions. The flow from the UPLC was coupled to a TimsTOF Flex (Bruker) instrument for mass spectrometric analysis, operated in both positive and negative ion mode. Compounds were annotated in Metaboscape (Bruker) using both an in-build rule-based annotation approach and a using LipidBlast MS2 library^91^. Features were removed if their average signal not were > 5 x more abundant in the QC samples than blanks (water extraction). The signals were normalized to internal standards in the SPLASH mix before correction for signal drift using statTarget^92^. Finally, signals were normalized using the QC samples^93^. Peak intensities from each measured metabolite were scaled using pareto scaling with *MetabolAnalyze* (version 1.3.1) and log_2_ transformed. We detected differentially abundant lipids using a linear model fitted separately for each lipid using limma^67^, using the model formula *E = 0 + timepoint + timepoint:exposure* which estimates each lipids difference in abundance between control and lps samples from the same time point. We generated all statistical values including *Q*-values using *ebayes*^67^.

### Statistics

Statistical methods are described above in respective sections. Analyses were made in R 4.02(https://www.r-project.org/) unless otherwise noted. Visualization was made using ggplot2^94^ unless otherwise noted.

## Acknowledgements

The authors would like to thank Navneet Vasistha, Diego García-González and all other members of the Khodosevich lab (BRIC, University of Copenhagen), Noor Irman, Michael Guldbrandsen, Eva Terrida (National Research Centre for the Working Environment, Copenhagen, Denmark), Christian Vaagensø, Jette Bornholdt (Department of Biology and BRIC, University of Copenhagen), Histocore (BRIC and Finsen laboratory, University of Copenhagen) for help with experiments, computational analysis, discussion and/or ideas.

## Author contributions

Conceptualization: ASa, KSH, SSKH. Supervision: ASa, KSH, KK. Experimental design: KSH, ASa, SSKH. Experimental work: SSKH, VA, BB, AP, JR, ACT, BE, NJF, JH, KSH. Data analysis and figure preparation: SSKH, DR, RK, VA, BB, NJS, JH, ASt, ASa, KSH. First paper draft: SSKH, ASa, KSH. Paper writing and editing: SSKH, RK, ASa, KSH, DR, UBV. Funding acquisition: ASa, KSH, SSKH.

## Data access

RNA-seq data has been deposited into the GEO database with accession number GSE224116

Mass spectrometry proteomics data have been deposited to the ProteomeXchange Consortium via the PRIDE partner repository. Data are available via ProteomeXchange with identifier PXD039402 [For referees: Username, Password: jRqf83O7]

Lipidomics raw data has been deposited at Figshare: https://doi.org/10.6084/m9.figshare.21967175.v1

## SUPPLEMENTARY METHODS

### Bronchoalveolar lavage

Lung inflammation in dams was assessed by collection of bronchoalveolar lavage fluid (BALf) followed by total and differential cell count^95^. Briefly, the trachea was cannulated by a 22 gauge needle equipped with a polyethylene catheter and the lungs were flushed twice by 0.8 ml of 0.9% NaCl through the trachea. Fluid was kept on ice until centrifugation at 400 × g at 4°C for 10 min. The pellet was resuspended in 100 μl medium (HAM F-12 with 1% penicillin/streptomycin and 10% fetal bovine serum). Aliquots of the cell suspension were used to determine numbers of live and dead cells by NucleoCounter (NC-200TM, Chemometec, Denmark), following manufacturer instructions. BALf cell composition (fractions of macrophages, lymphocytes, neutrophils and epithelial cells) was determined following centrifugation of 40 μl of suspension at 55xg for 4 min (Cytofuge 2, Statspin, TRIOLAB, Brøndby, DAnmark) on to a microscope slide followed by fixation in 96% ethanol and staining with May-Grünwald-Giemsa. A minimum of 200 cells/slide were counted under a light microscope. All slides were randomized, blinded and scored by the same technician on the same day. By combining differential cell counts with cell numbers, the number of cells in each fraction could be calculated.

### Saa3 RT-PCR analysis

We quantified lung mRNA expression levels of the acute phase response gene *Saa3*. RNA was isolated from 16–20 mg of tissue on Maxwell® 16 (Promega, USA) using Maxwell® 16 LEV simply RNA Tissue Kit (AS1280, Promega, USA) according to the manufacturer’s protocol. RNA was eluted in 50 μl nuclease free (DEPC) water. cDNA was prepared from DNase treated RNA using Taq-Man® reverse transcription reagents (Applied Biosystems, USA) following the manufacturer’s protocol. Total RNA and cDNA concentrations were measured on NanoDrop 2000c (ThermoFisher, USA). The Saa3 mRNA levels were determined using real-time RT-PCR with 18S RNA as reference gene. Each sample was run in triplicates on the ViiA7 Real-Time PCR (Applied Biosystems, USA). Saa3 primers and probe sequences were: forward: 5’ GCC TGG GCT GCT AAA GTC AT 3′, reverse: 5’ TGC TCC ATG TCC CGT GAA C 3′ and Saa3 probe: 5’ FAM-TCT GAA CAG CCT CTC TGG CAT CGC T-TAMRA 3′. In all assays, TaqMan pre-developed mastermix (Applied Biosystems, USA) was used. Saa3 and 18S RNA levels were quantified in triplicates in separate wells. The relative expression levels of the target gene were calculated by the comparative method 2-ΔCt. Negative controls, where RNA had not been converted to cDNA (no template control), were included in each run. One sample, the plate control, was included in all Real-Time PCR analyses.

## SUPPLEMENTARY NOTES

### Note S1: Choice of LPS dose

LPS dose was chosen to model robust airway inflammation without causing excessive lung injury or preterm birth. In a pilot study, we administered 1.5, 4 or 7 ug LPS/animal or vehicle to non-pregnant mice. The lowest dose caused ∼5% weight-loss after 24h (Fig. S1A) and was thus within the upper range of accepted body weight loss (10%) in short-term toxicity studies^96^, indicative of maximum tolerable dose. At the two higher doses weight loss exceeded 10% after 24h. Pulmonary neutrophil-influx (a proxy for pulmonary inflammation) was similar across the three dosages after 24h, indicating saturation of neutrophil-influx already at 1.5 µg (Fig. S1B). As we were interested in placental transfer of inflammation and not outright toxicity, we chose the dose of 1 µg LPS/animal for our study.

### Note S2: Development genes downregulated in fetal liver

At 5h, a number of genes associated with GO terms related to development were downregulated at 5h, including mesenchyme development, cell differentiation and ventricular and cardiac septum development (Fig.S4A). Of note, there is an overlap in genes responsible for liver and heart development. On gene level, we observed transient downregulation of growth factors and their positive regulators (e.g. *Smad4, Cited2, Ctnnb1, Fgfr2, Dand5*) and transcription factors (*Sox9, Lef1, Snai1/2*), mostly confined to 5h.

### Note S3: Glycosylation genes downregulated in fetal liver

Genes associated with glycosylation-related GO-terms (Fig.S4B) were downregulated at 5h. Interestingly, some of these genes overlapped the downregulated glycosylation-associated genes in the placenta at 12h, including the *Dag1* associated genes *Pomgnt1-2*, *Pomt1-2* and *Fkrp*, and the Notch signaling-associated genes *Pofut1* and *Poglut1*. Additional dystroglycan and Notch-pathway associated genes were downregulated in fetal liver only, including *B3galnt2*, *Pomk* and *Poglut2*. Since dystroglycan glycosylation encourages a loosening in cell-cell adhesion and tissue permeability, we suggest that the dystroglycan decrease in fetal liver increases permeability and nutrient uptake during a state of scarcity. A group of genes related to glycosylation of glucose and carbohydrates (e.g., *B3gat3*, *Poglut2*, *B3galt6*, *Poglut3*, *St6gal1*) were also downregulated. Alterations in glycosylation profiles of proteins involved in lipoprotein metabolism are associated with changes in their function^97^ so we suggest that the alterations in carbohydrate and monosaccharide intermediates may also be an effect of the decreased cholesterol and fatty acid metabolism.

## SUPPLEMENTARY TABLE LEGENDS

**Table S1: GO term analysis**

GO term over-representation analysis for all tissues and time points. Available at https://doi.org/10.6084/m9.figshare.22059308.v1

## SUPPLEMENTARY FIGURES WITH LEGENDS

**Figure S1.**
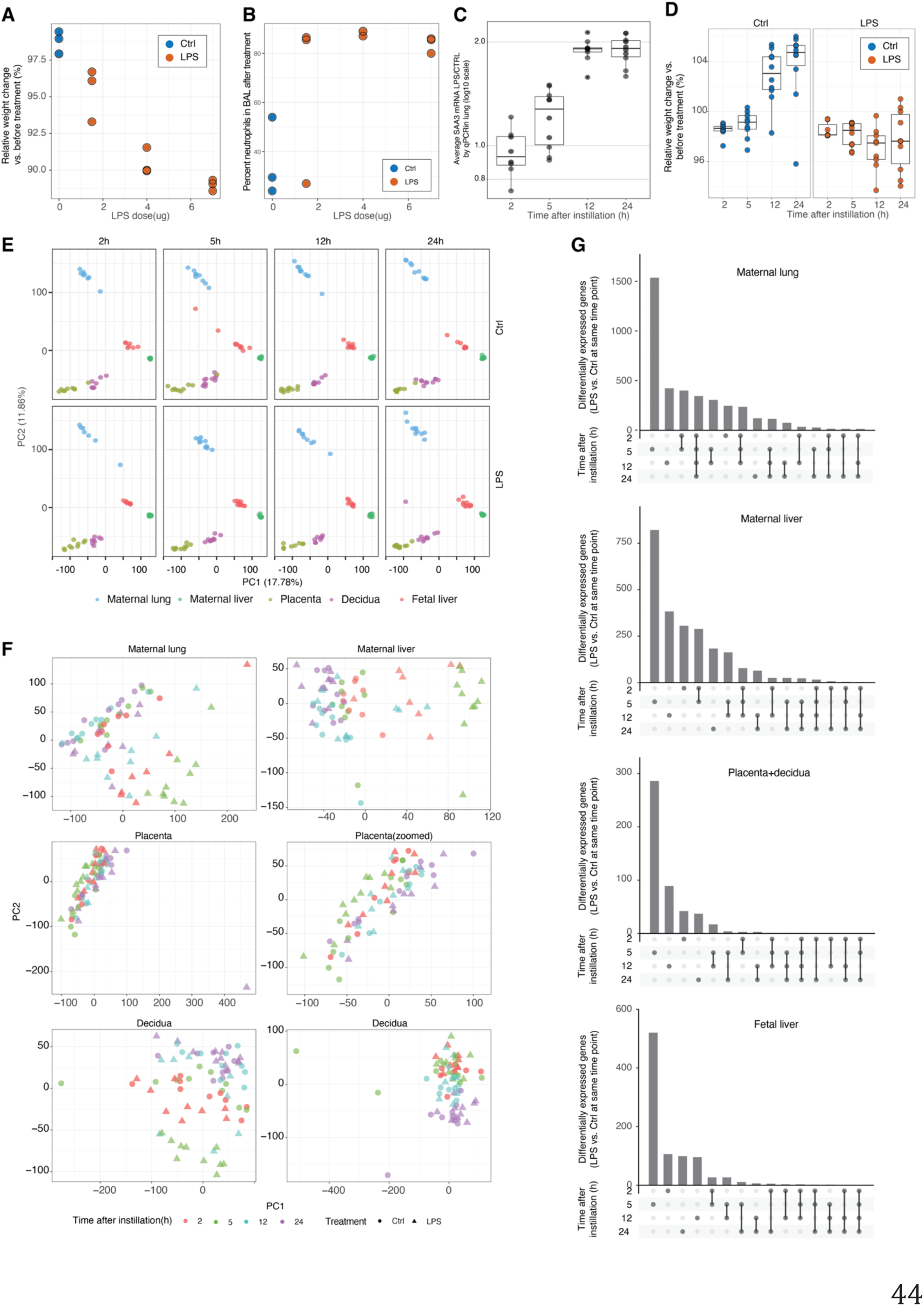
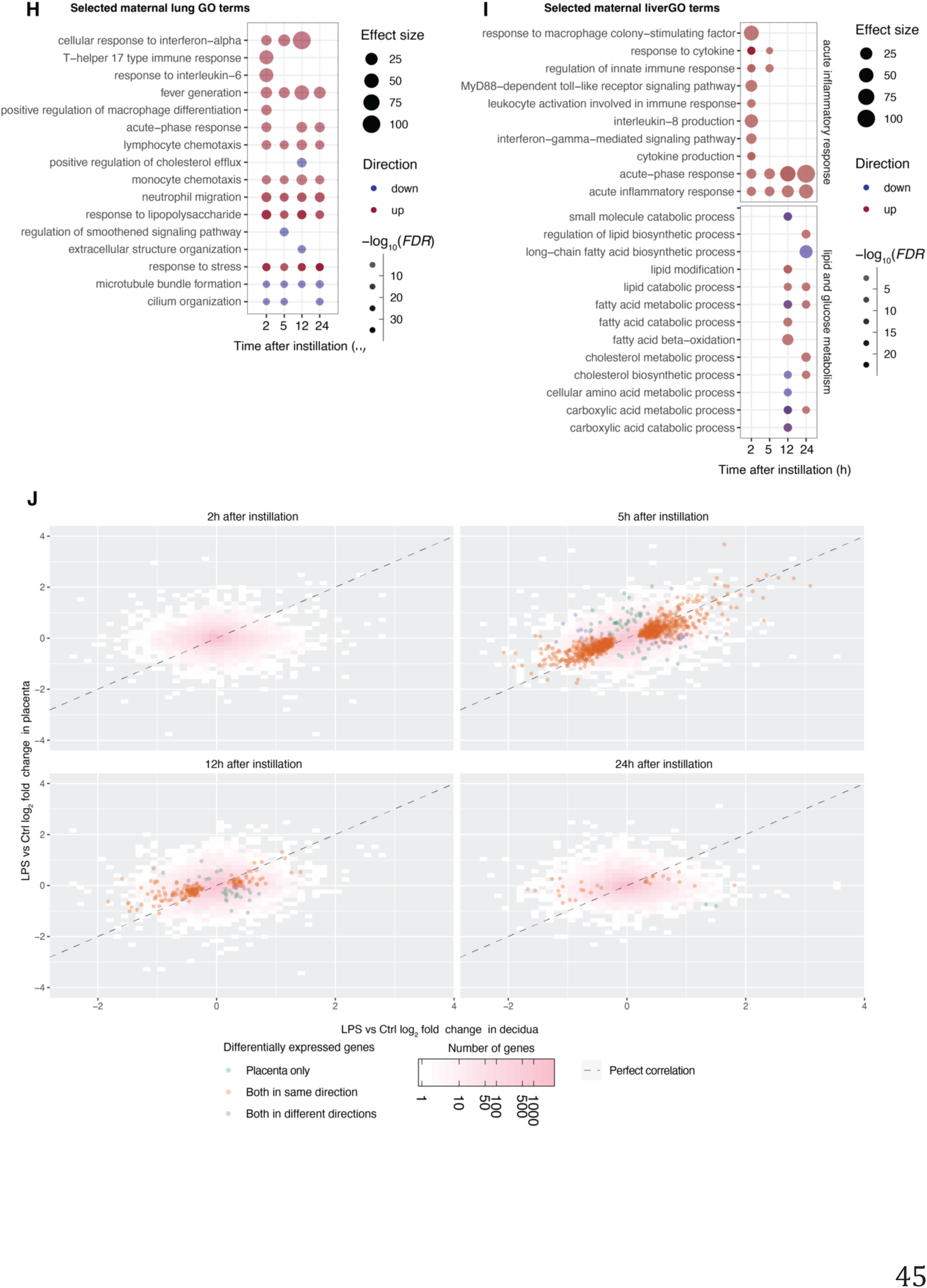
This figure expands figure 1. **A: Weight loss as a function of LPS dose (pilot study).** The same type of LPS intratracheal instillation as in the main study was performed, but with varying LPS doses. Y axis shows relative weight (%) of mice compared to before treatment as a function of LPS dose (X axis). Dots indicate individual mice, colored by treatment. Mice were weighed before instillation and before euthanization (24h after LPS-exposure). **B: Neutrophils in BAL as a function of LPS dose (pilot study).** Neutrophil % in BAL were measured in the same mice as in A. The Y axis shows % neutrophils in BAL. The X axis shows LPS dose as in A. Dots indicate individual mice, colored by treatment. Samples were collected 24h after LPS exposure. **C: Expression change of Saa3 following LPS treatment.** *Saa3* gene expression was measured in mice lungs after instillation with LPS or Ctrl by real-time RTPCR. Dots on the Y axis shows each LPS-instilled lung *Saa3* expression value divided by the average Ctrl *Saa3* expression, on log_10_ scale. **D: Relative weight change of mice in the main experiment.** Y axis shows % change compared to before treatment (Ctrl or LPS). Box plots and individual mice (dots) are shown, colored by treatment. X axis shows time after instillation. **E: Principal component analysis (PCA) of RNA-seq data, organized by time after instillation and treatment.** X axis shows PC1, Y axis shows PC2 (% variance explained per PC is shown). Panel columns indicate time after instillation. Panel rows indicate treatment. Dots indicate individual dams or fetuses, colored by tissue. **F: PCA of RNA-seq data, organized by tissue.** Each panel shows RNA-seq data from one tissue; for the placenta, a zoom-in view is also shown due to one outlier sample. X axis shows PC1, Y axis shows PC2. Dots indicate individual mice. Dot color shows time after instillation. Dot shape shows treatment. **G: UpSet plot visualization of number of differentially expressed (DE) genes and their overlap across time.** Each upset plot shows the results for differential expression analysis in one tissue, as indicated on top. The top barplot in each upset plot shows the number of DE genes (LPS vs Ctrl, *FDR*<0.05) for one tissue on the Y axis. Time points are shown as rows below the barplot. Dots indicate what overlap of time point(s) that is plotted in the bar plot above. Lines between dots indicate genes DE across two or more time points. **H: Gene ontology analysis of maternal lung RNA-seq data**. X axis shows time after instillation. Rows show selected GO terms (see Table S1 for all terms). Dots show term over-representation in up- (red) or down-regulated (blue) genes following LPS instillation. Dot size and color intensity show enrichment effect size and statistical significance (-log_10_(*FDR*)) **I: Gene ontology analysis of maternal liver RNA-seq data**. Plot is organized as in H, but shows analysis of maternal liver RNA-seq data, and splits GO terms into two larger families, as indicated to the right. **J: Correlation between LPS-instillation response between placenta and decidua**. In each subplot, Y axis shows the LPS vs Ctrl log_2_FC expression change values of genes based on RNA-seq data from the placenta, while X axis shows corresponding log_2_FC values from decidua. Rectangle color shows the number of genes in each position of the plot. Dots show individual genes that are DE (LPS vs Ctrl, *FDR*<0.05) in placenta only, in both tissues in the same direction, and in both tissues but in opposite directions, as indicated by color. Each subplot shows one time point after instillation, as indicated on top. Dotted lines show perfect correlation (X=Y) lines.

**Fig. S2:**
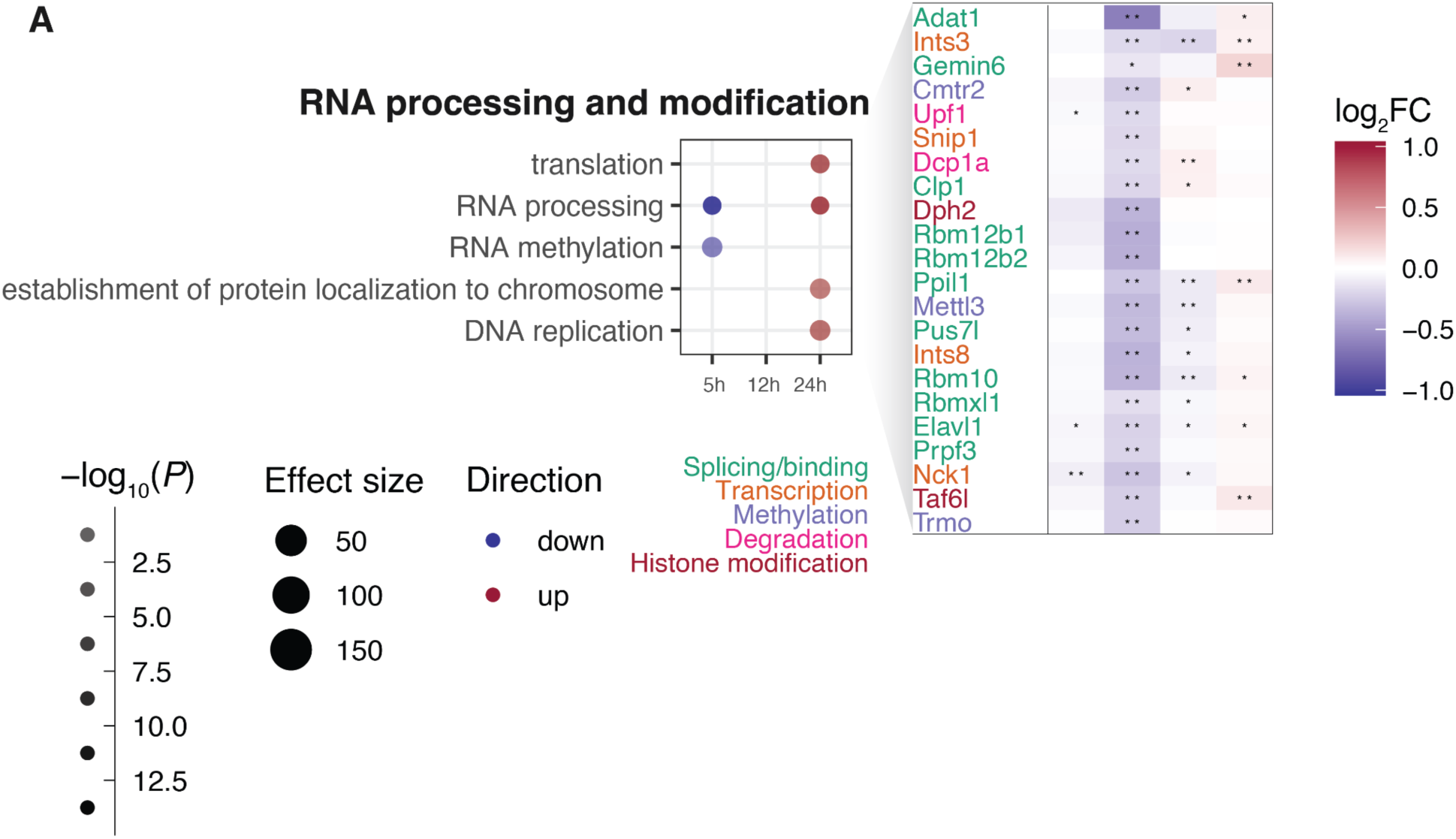
Detailed analysis of placental gene expression change. This figure expands Fig.3 with an additional GO theme and associated genes, organized in the same way as the left and middle panels of Fig.3.

**Fig. S3:**
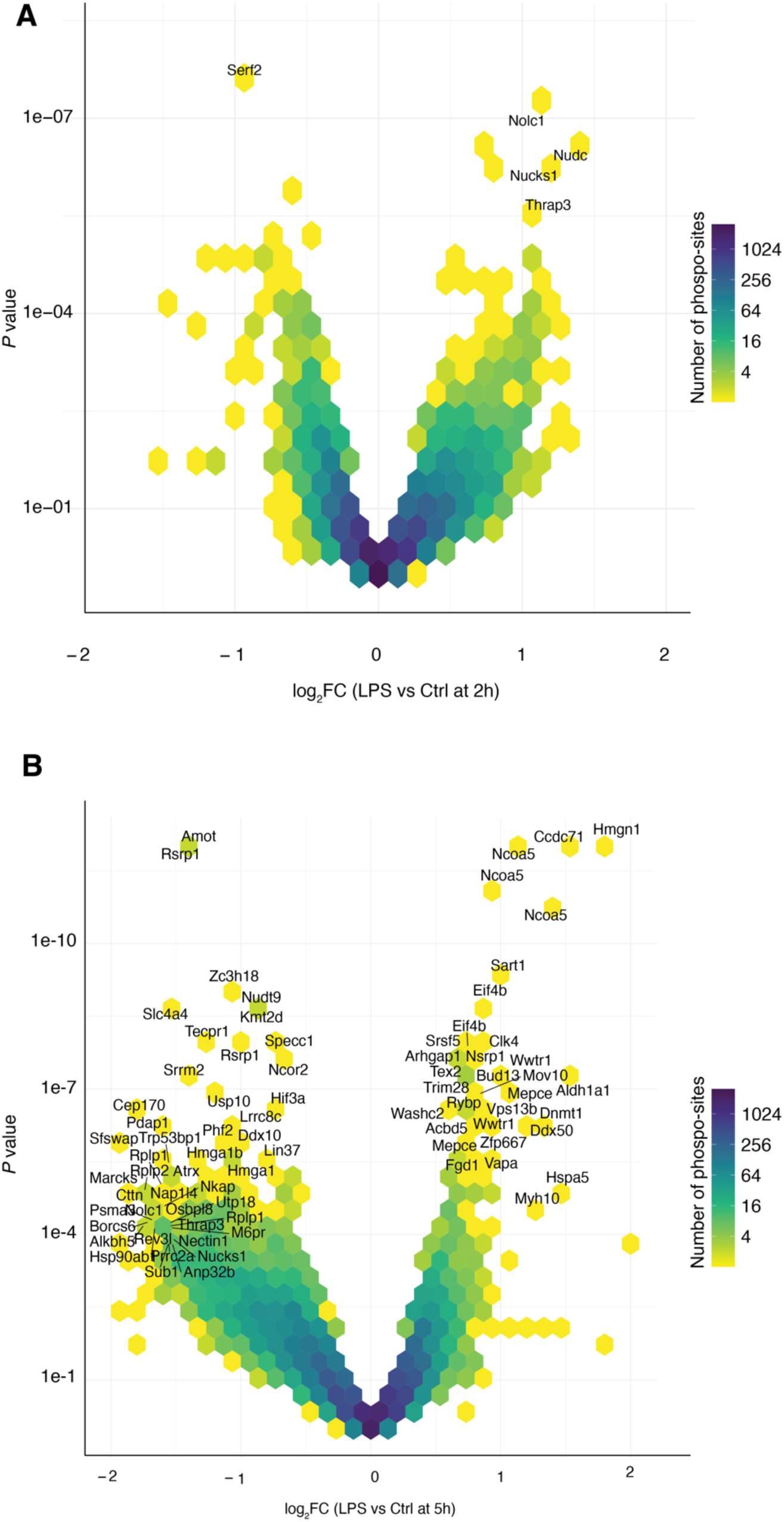
Phospoproteomics analysis of early placental time points. This figure expands Fig.4. **A: Phosphorylation site changes at 2h in placenta.** The X axis shows LPS vs Ctrl log_2_FC based on phosphoproteomics analysis (negative values correspond to loss of phosphorylation, positive to gain). The Y axis shows corresponding *P* values using −log_10_ scale. Hexagon colors show the number of phospho-sites in a given region in the plot. Genes with substantial phosphosites are labeled, where genes mentioned in the main text are in bold. **B: Phosphorylation site changes at 5h in placenta.** Plot is organized as in panel A, but shows data from 5h.

**Fig. S4:**
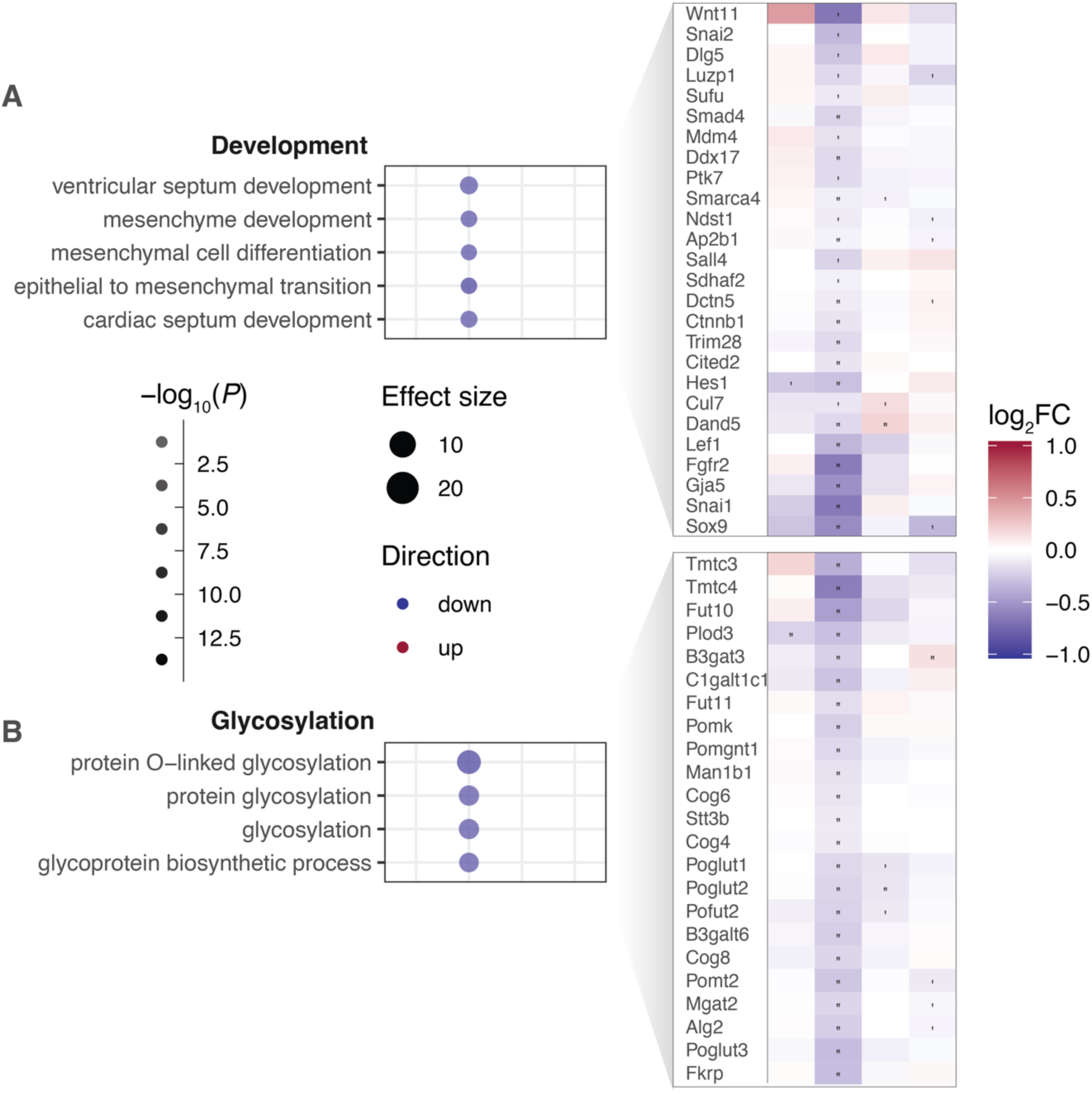
Detailed analysis of fetal liver gene expression change. This figure expands Fig.5 with two additional GO themes and associated genes, organized in the same way as the left and middle panels of Fig.4.

**Fig. S5:**
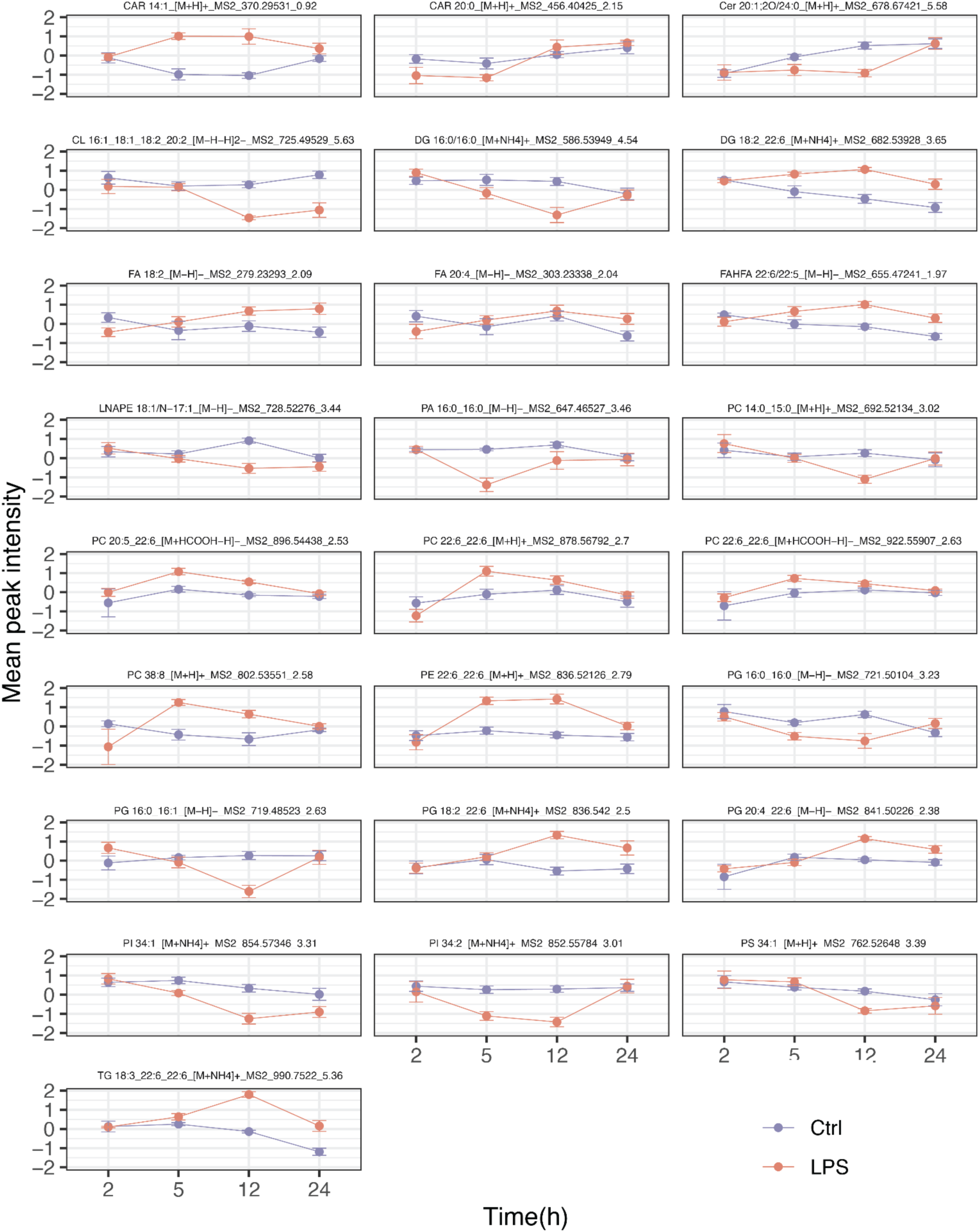
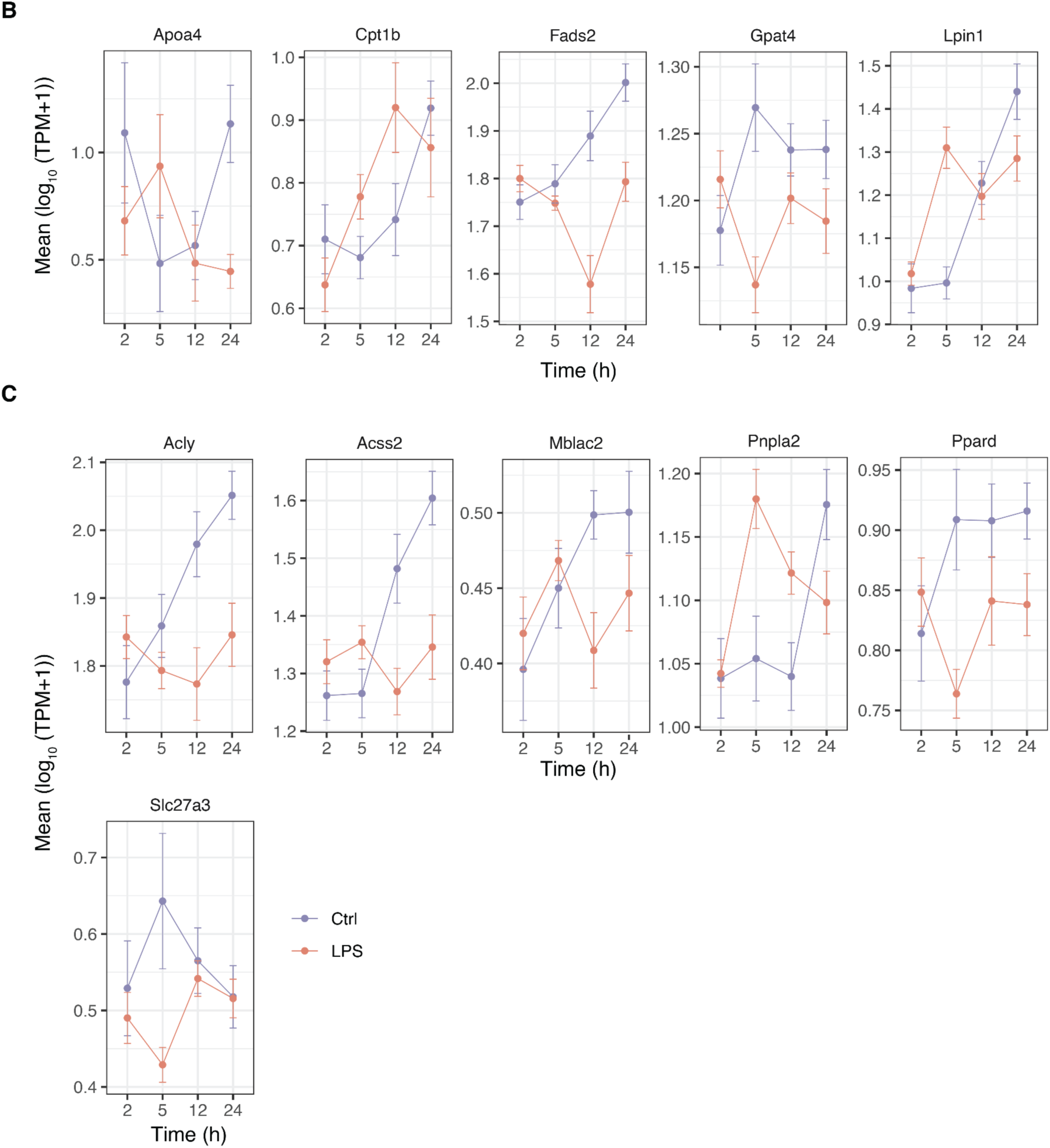
Lipid- and associated RNA abundance related to synthesis of DHA-containing lipid species. This figure expands Fig.6. A: Average lipid abundance for all lipids in **Fig 6C**. Images are organized as in the callout figures in Fig.6C. B: Average mRNA expression of genes in **Fig 6C**. Images are organized as in the callout figures in Fig.6C, but the Y axis shows log_10_(average RNA-seq TPM+1); gene names are shown on top. C: Average mRNA expression of genes using AcetylCoA as a substrate in fetal liver. Images are organized as in panel B.

